# Conservation and divergence of UVR8-COP1/SPA-HY5 signaling in UV-B responses of *Marchantia polymorpha*

**DOI:** 10.1101/2025.07.16.665153

**Authors:** Yuanke Liang, Roman Podolec, Richard Chappuis, Emmanuel Defossez, Gaétan Glauser, Johannes Rötzer, Sara Christina Stolze, Liam Dolan, Hirofumi Nakagami, Emilie Demarsy, Roman Ulm

## Abstract

Ultraviolet-B radiation (UV-B) poses a major challenge to all forms of plant life. The liverwort *Marchantia polymorpha* (Marchantia) serves as a key model organism to study signaling pathways and to infer their evolution throughout the green lineage. Marchantia expresses key components of UV-B signaling, including the photoreceptor UV RESISTANCE LOCUS 8 (MpUVR8), the WD40-repeat protein REPRESSOR OF UV-B PHOTOMORPHOGENESIS (MpRUP), the E3 ubiquitin ligase complex CONSTITUTIVELY PHOTOMORPHOGENIC 1 / SUPPRESSOR OF phyA-105 (MpCOP1/MpSPA), and the transcriptional regulator ELONGATED HYPOCOTYL 5 (MpHY5). Here, we show that MpUVR8 exists as a homodimer in its ground-state in vivo, then monomerizes and accumulates in the nucleus upon UV-B activation. Activated MpUVR8 interacts with MpCOP1, triggering growth inhibition, genome-wide gene expression changes, biosynthesis of UV-absorbing metabolites, and photoprotection, which overall contributes to UV-B stress tolerance. MpRUP facilitates redimerization of MpUVR8 and Mp*rup* null mutants show enhanced UV-B photomorphogenesis, demonstrating that MpRUP efficiently represses MpUVR8 signaling. Unlike the case in Arabidopsis and in contrast to the strong Mp*cop1* mutant phenotype, Mp*spa* mutants develop only a very weak constitutive photomorphogenesis phenotype, indicating that COP1 function is much more independent of SPA in Marchantia than in Arabidopsis. Moreover, in contrast to Arabidopsis SPAs, Mp*spa* is linked with a hyper-responsive UV-B phenotype, suggesting that MpSPA is a negative regulator of MpUVR8 signaling. Similar to Arabidopsis HY5/HYH, MpHY5 functions antagonistically to MpCOP1, but its role in UV-B-mediated gene expression changes is more limited. Our findings demonstrate that although core components of UV-B signaling existed in the last common ancestor of extant land plants, regulatory interactions have diversified in different lineages since their divergence more than 400 million years ago.

## INTRODUCTION

Photoautotrophic plants have a “love/hate relationship” with light. Light is required for photosynthesis, but sunlight poses major challenges due to potentially damaging ultraviolet-B radiation (UV-B; 280–315 nm) (Demarsy et al., 2018). Plants have a unique and conserved UV-B photoreceptor UV RESISTANCE LOCUS 8 (UVR8) that is crucially important for UV-B acclimation, tolerance, and survival (Kliebenstein et al., 2002; Rizzini et al., 2011; Tilbrook et al., 2016; Li et al., 2018; Soriano et al., 2018; Rai et al., 2019; Podolec et al., 2021a; Hu et al., 2024; Stockenhuber et al., 2024). The UVR8 signaling pathway has mainly been outlined in Arabidopsis, with UVR8 being a homodimer in its ground state, which exhibits monomerization after absorption of UV-B photons by intrinsic tryptophan (Trp) residues, followed by accumulation in the nucleus (Kaiserli and Jenkins, 2007; Rizzini et al., 2011; Christie et al., 2012; Wu et al., 2012; Yin et al., 2016; Li et al., 2020; Qian et al., 2020; Podolec et al., 2021a; Fang et al., 2022). Activated UVR8 then interacts with CONSTITUTIVELY PHOTOMORPHOGENIC 1 (COP1) (Favory et al., 2009; Yin et al., 2015; Wang et al., 2022). COP1 forms an E3 ubiquitin ligase complex together with SUPPRESSOR OF phyA-105 proteins (SPA1–4 in Arabidopsis) (Laubinger et al., 2004; Chen et al., 2010). The COP1/SPA complex represses photomorphogenesis by ubiquitinating photomorphogenesis-promoting factors, directing them to degradation by the 26S proteasome (Podolec and Ulm, 2018; Ponnu and Hoecker, 2021). UV-B-activated UVR8 inhibits the activity of the COP1-SPA complex through a cooperative, high-affinity binding mechanism that displaces substrate proteins from COP1, thus allowing the stabilization of key transcription factors, including the basic leucine zipper (bZIP) transcriptional regulators ELONGATED HYPOCOTYL 5 (HY5) and HY5 HOMOLOG (HYH) (Holm et al., 2002; Ulm et al., 2004; Oravecz et al., 2006; Binkert et al., 2014; Lau et al., 2019). This ultimately leads to broad changes in gene expression upon UV-B exposure (Brown et al., 2005; Favory et al., 2009; Tavridou et al., 2020b). COP1/SPA together with HY5/HYH thus represents a major regulatory hub in light signaling in Arabidopsis (Holm et al., 2002; Ulm et al., 2004; Ponnu et al., 2019). The two WD-40 repeat proteins REPRESSOR OF UV-B PHOTOMORPHOGENESIS 1 (RUP1) and RUP2 act as negative feedback regulators of the UVR8 photocycle by facilitating UVR8 redimerization, thereby repressing UV-B signaling (Gruber et al., 2010; Heijde and Ulm, 2013; Podolec et al., 2021b; Wang et al., 2023).

The core components of UVR8 signaling seem to be evolutionarily conserved in the green lineage, since they are present in the alga *Chlamydomonas reinhardtii* (hereafter Chlamydomonas) (Allorent et al., 2016; Tilbrook et al., 2016; Tokutsu et al., 2019; Dannay et al., 2025) and the liverwort *Marchantia polymorpha* (hereafter Marchantia) (Clayton et al., 2018; Soriano et al., 2018; Kondou et al., 2019). Notably, Marchantia has minimal gene redundancy in many signaling pathways, and many studies in the past decade have contributed to our understanding of the origins and evolution of major plant regulatory pathways (Bowman et al., 2017; Liang et al., 2022; Frangedakis et al., 2024). The Marchantia genome indeed possesses single homologs for genes encoding UVR8 signaling-related components, such as MpUVR8, MpRUP, MpCOP1, MpSPA, and MpHY5. Although a few studies have explored the role of some of these components in UV-B (Clayton et al., 2018; Soriano et al., 2018; Kondou et al., 2019) and visible light signaling (Zhang et al., 2021; Yelina et al., 2024), a deeper understanding of their activities and roles in UV-acclimation is warranted to understand to what extent the molecular mechanisms by which these components mediate UV acclimation and tolerance are conserved or divergent throughout the plant kingdom.

We investigated if the major UVR8 signaling components have functionally diverged during plant evolution, and how mechanisms underlying UV-B tolerance may have evolved during the colonization of land by plants. We extend our understanding of the crucial role of MpUVR8 in UV-B acclimation and survival in Marchantia, and provide unequivocal evidence clarifying conservation of the UVR8 photocycle in Marchantia. However, although Mp*cop1* mutants exhibited enhanced constitutive photomorphogenesis and extreme dwarfism comparable to UV-B-grown Mp*rup* mutants, Mp*spa* mutants showed only a very weak constitutive photomorphogenesis phenotype in visible light. Surprisingly however, Mp*spa* mutants were hyper-responsive to UV-B, suggesting that MpSPA plays a divergent role from that in Arabidopsis. We further provide a genome-wide view of MpUVR8-mediated transcriptional responses in Marchantia and identify UV-B-regulated gene clusters and metabolites key to UV-B protection. Notably, different to HY5/HYH being major regulators in Arabidopsis, our transcriptome data revealed that only a few UV-B-regulated genes were entirely MpHY5-dependent.

## RESULTS

### MpUVR8 is homodimeric in its ground state, and monomerizes and accumulates in nuclei in response to UV-B

In Marchantia, UVR8 is encoded by a single locus, but alternative splicing of the 11^th^ intron produces two transcripts, namely Mp*UVR8.1* and Mp*UVR8.2*, resulting in two different proteins whereby residues 414 to 426 present in MpUVR8.1 are absent in MpUVR8.2 (fig. S1) (Soriano et al., 2018). Whether MpUVR8 forms homodimers *in vivo* or is primarily monomeric with a divergent inactivation mechanism is unclear (Soriano et al., 2018). We thus first tested if MpUVR8.1 and MpUVR8.2 form dimers using a luciferase complementation imaging (LCI) assay in transiently transformed *Nicotiana benthamiana* leaves (Zhou et al., 2018). Resulting LCI signals showed interactions of MpUVR8.1 and MpUVR8.2 both with and between themselves, indicating that MpUVR8 forms homodimers *in vivo* (Fig. 1A to C). In AtUVR8, the aspartate-96 (D96) and aspartate-107 (D107) residues are key to homodimer formation and mutating each to asparagine impairs UVR8^D96N,D107N^ dimer formation (Christie et al., 2012; Wu et al., 2012; Heilmann et al., 2016; Podolec et al., 2021b). Consistent with the conservation of these residues in MpUVR8 (fig. S1), no LCI signal was detected with the mutated MpUVR8.1^D95N,D106N^ and MpUVR8.2^D95N,D106N^ variants (Fig. 1A and B).

**Figure 1.**
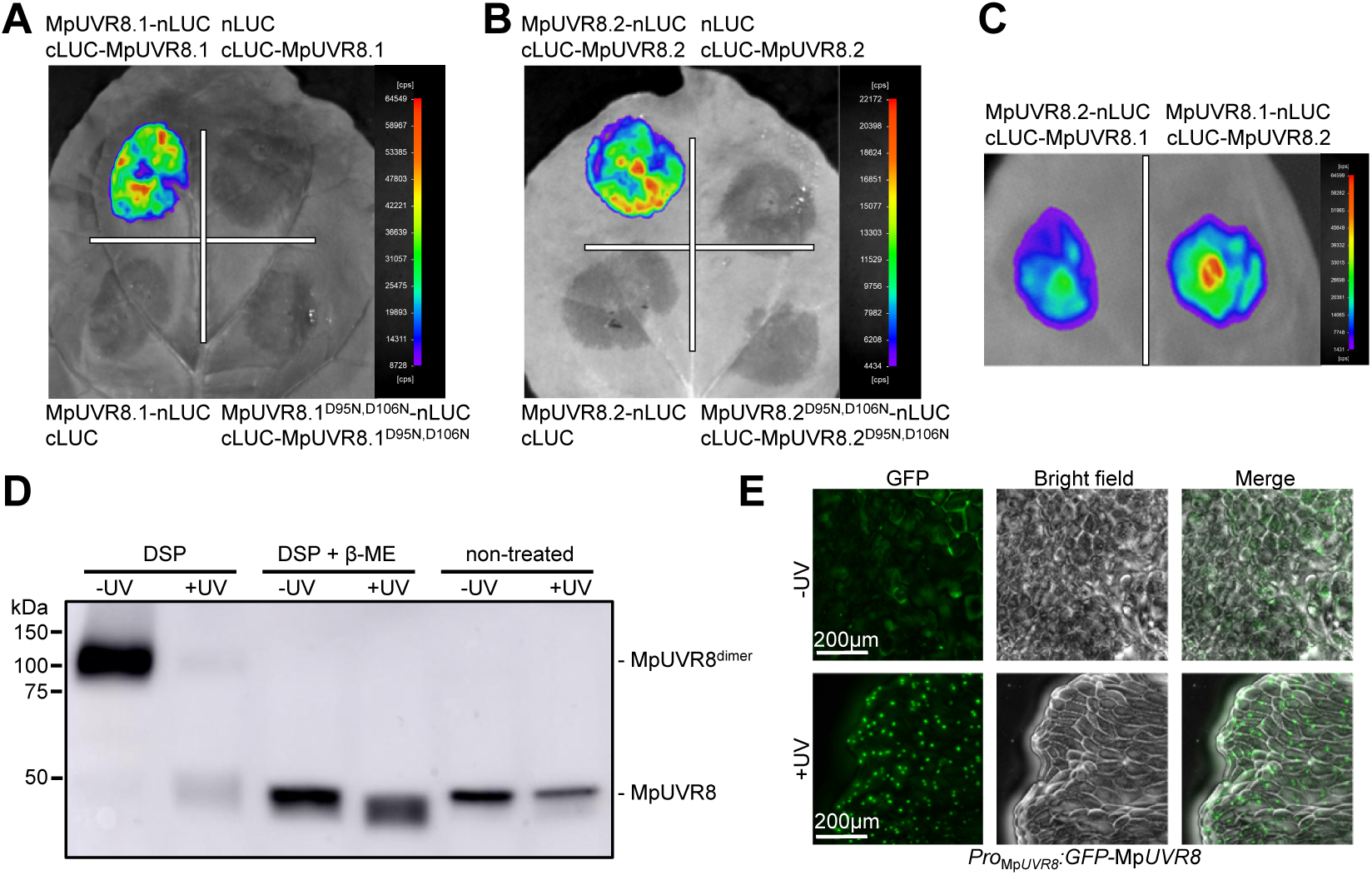
MpUVR8.1 and MpUVR8.2 form dimers in their ground state, monomerize upon UV-B exposure, and accumulate in the nucleus. **A-C)** Luciferase complementation imaging (LCI) assays assessing monomer interaction for MpUVR8.1 (A) and MpUVR8.2 (B), as well as their D95N,D106N variants, and MpUVR8.1 and MpUVR8.2 (C). MpUVR8 versions were fused to N-terminal (nLUC) or C-terminal (cLUC) fragments of luciferase, expressed in *N. benthamiana* leaves and imaged 2 days after infiltration. An image of a representative leaf is shown. Cps, counts per second. **D)** Immunoblot analysis of MpUVR8 dimer/monomer levels in 7-day-old wild type (Tak-1) gemmalings irradiated (+UV) or not (-UV) for 20 min with broadband UV-B. Protein extracts were either crosslinked with DSP, DSP crosslinked then reversed with the reducing agent β-mercaptoethanol (DSP + β-ME), or non-treated. **E)** Subcellular localization of GFP-MpUVR8 in 7-day-old gemmalings of a Tak-1/*Pro*_Mp*UVR8*_*:GFP-*Mp*UVR8* line, grown in white light and irradiated or not for 20 min with 25 μmol m^-2^ s^-1^ broadband UV-B (+UV and -UV, respectively). Bars = 200 μm.

To further test if endogenous MpUVR8 forms homodimers, we crosslinked protein extracts from Marchantia with dithiobis (succinimidyl propionate) (DSP), as described previously for Arabidopsis (Rizzini et al., 2011; Podolec et al., 2021b). Using a specific anti-MpUVR8 antibody, we detected a band at the size of the predicted MpUVR8 dimer at about 100 kDa in the absence of UV-B, and this band was strongly reduced in extracts from UV-B-treated plants (Fig. 1D). These data indicate that UV-B exposure induces MpUVR8 monomerization. Of note, UV-B-activated monomeric MpUVR8 was not well detected in crosslinked extracts, as in Arabidopsis (Rizzini et al., 2011; Podolec et al., 2021b). However, we detected MpUVR8 at its monomeric size of about 48 kDa (predicted sizes for MpUVR8.1 and MpUVR8.2 are 48.6 and 47.1 kDa, respectively) both following the reversal of DSP crosslinking with the reducing agent β-mercaptoethanol (β-ME) and in the absence of crosslinking (Fig. 1D), demonstrating that MpUVR8 levels are not affected by UV-B treatment or crosslinking.

To define MpUVR8 subcellular localization in Marchantia, we transformed wild-type Tak-1 with the full genomic Mp*UVR8* locus containing an N-terminal green fluorescent protein (GFP) tag and imaged the resulting fluorescence. GFP-MpUVR8 accumulated in the nucleus only when plants were exposed to UV-B (Fig. 1E), similar to AtUVR8 (Kaiserli and Jenkins, 2007; Yin et al., 2016) and the 35S promoter–driven expression of Citrine-MpUVR8 in Tak-1 (Kondou et al., 2019). This contrasts with the apparent constitutive nuclear localization of GFP-MpUVR8.1 and GFP-MpUVR8.2 when expressed in Arabidopsis and *N. benthamiana* (Soriano et al., 2018). We conclude that upon UV-B exposure MpUVR8 monomerize and accumulate in nuclei in Marchantia.

### MpRUP facilitates MpUVR8 redimerization

We tested if MpUVR8 interacts with MpRUP, the putative ortholog of AtRUP1 and AtRUP2 (Gruber et al., 2010; Clayton et al., 2018). MpUVR8 and MpRUP exhibited interaction in yeast two-hybrid (Fig. 2A and B) and LCI assays (Fig. 2C, D), and MpRUP co-purified with GFP-MpUVR8 in affinity-purification coupled with mass spectrometry (AP-MS) (fig. S2). We further assayed if MpRUP facilitates MpUVR8 redimerization in Marchantia. In wild-type Tak-1 plants, MpUVR8 dimer amount was strongly reduced upon exposure to UV-B, then reverted to initial levels within approximately 1 h once under -UV-B conditions (Fig. 2E). However, in a Mp*rup* mutant generated using CRISPR/Cas9 mutagenesis in the Tak-1 background (fig. S3), MpUVR8 dimer levels recovered slowly and did not reach “-UV” ground state levels even after 6 h (Fig. 2E). We thus conclude that MpRUP facilitates MpUVR8 redimerization, indicating a conserved mechanism for negative feedback regulation of UVR8 activity.

**Figure 2.**
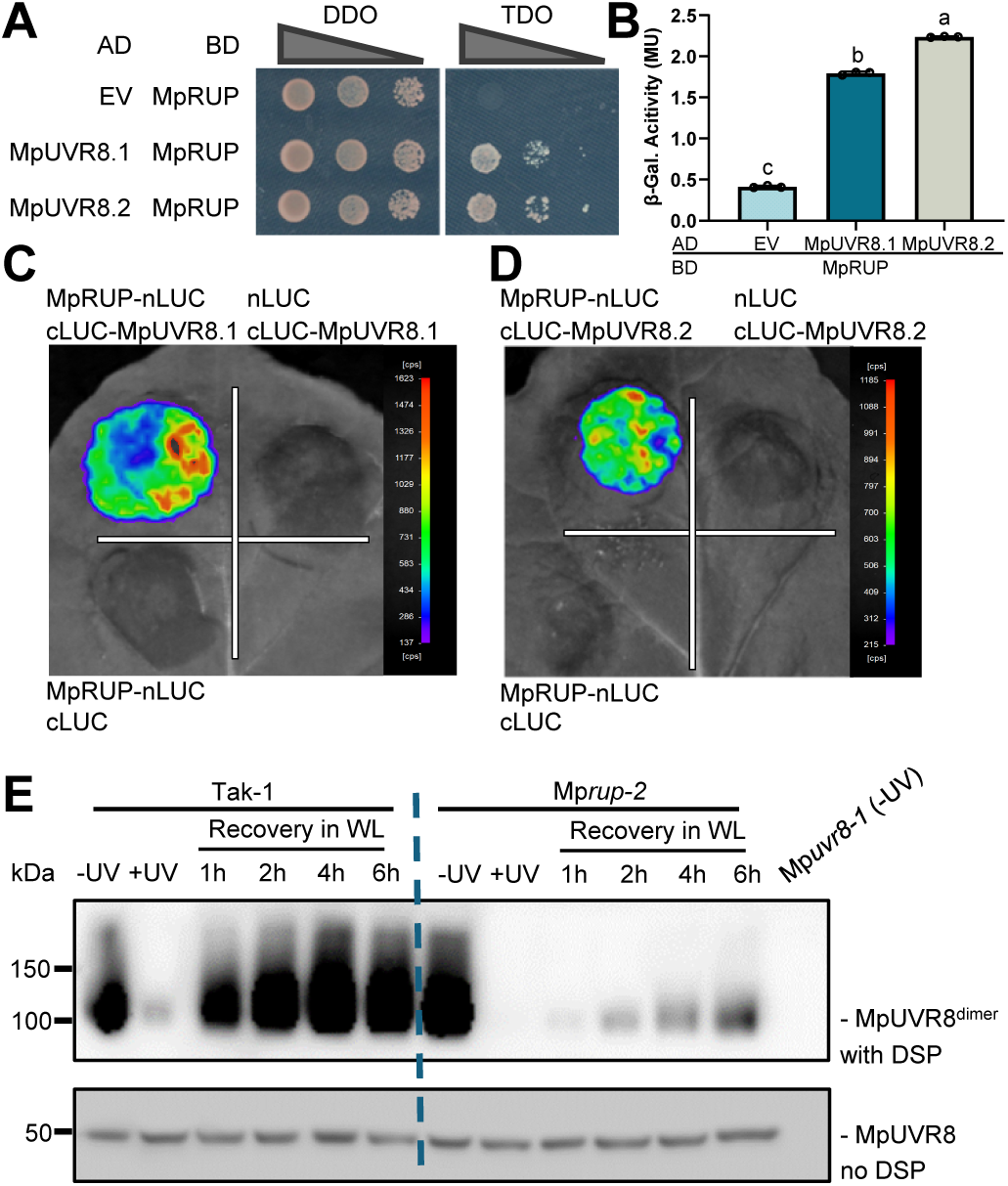
MpRUP facilitates MpUVR8 redimerization. **A)** Yeast two-hybrid assays of MpUVR8.1 and MpUVR8.2 interactions with MpRUP. Representative images of serial dilutions spotted on non-selective SD/-Trp/-Leu double dropout (DDO) and selective SD/-Trp/-Leu/-His triple dropout (TDO) media are shown. AD, transcriptional activation domain; BD, DNA-binding domain; EV, empty vector. **B)** Quantitative yeast two-hybrid assays of MpUVR8.1 and MpUVR8.2 interactions with MpRUP. Individual data points of biological replicates and means ± SD are shown (*n* = 3). Different letters indicate statistically significant differences between the means (P < 0.01) as determined by one-way ANOVA followed by Tukey’s post hoc honestly significant difference (HSD) test. β-Gal., β-galactosidase; MU, Miller units. **C** and **D)** LCI assays of MpRUP interactions with MpUVR8.1 (C) and MpUVR8.2 (D) fused to N-terminal (nLUC) or C-terminal (cLUC) luciferase fragments, expressed in *N. benthamiana* leaves and imaged 2 days after infiltration. Images of representative leaves are shown. Cps, counts per second. **E)** Immunoblot analysis of MpUVR8 levels in 7-day-old wild type (Tak-1) and Mp*rup-2* gemmalings irradiated for 20 min with broadband UV-B (+UV) before recovery in white light (WL) for the indicated times (1–6 h). Non-irradiated samples (-UV) are included as MpUVR8 ground state controls alongside a negative control -UV Mp*uvr8-1* sample. Top: MpUVR8 dimer levels in DSP-crosslinked protein extracts; bottom: total MpUVR8 levels in non-crosslinked extracts.

### MpUVR8 and MpRUP antagonistically regulate UV-B-dependent growth, pigment accumulation, and acclimation

To explore the phenotypic consequences of MpUVR8 and MpRUP action, we established robust UV-B conditions for phenotypic and molecular analysis. Under continuous white light, no striking differences were detectable between Tak-1, Mp*uvr8*, and Mp*rup* (Fig. 3A and fig. S4). However, when exposed to white light supplemented with narrowband UV-B (1.5 μmol m^-2^ s^-1^), Tak-1 exhibited a slight growth inhibition and slight accumulation of pigments (Fig. 3A and fig. S4A). These responses were reduced, if not absent, in Mp*uvr8* mutant and Mp*RUP^OE^* overexpression lines (Fig. 3A and fig. S4A, F). By sharp contrast, Mp*rup* mutants exhibited a striking UV-B-hyperresponsive photomorphogenic phenotype, with extreme growth inhibition and strong pigmentation of the thallus specifically under UV-B (Fig. 3A and fig. S4A) (Clayton et al., 2018), similar to transgenic lines overexpressing Mp*UVR8.1* or Mp*UVR8.2* photoreceptors (fig. S4A, E). We conclude that MpRUP plays a major role in MpUVR8 inactivation to sustain growth under UV-B.

**Figure 3.**
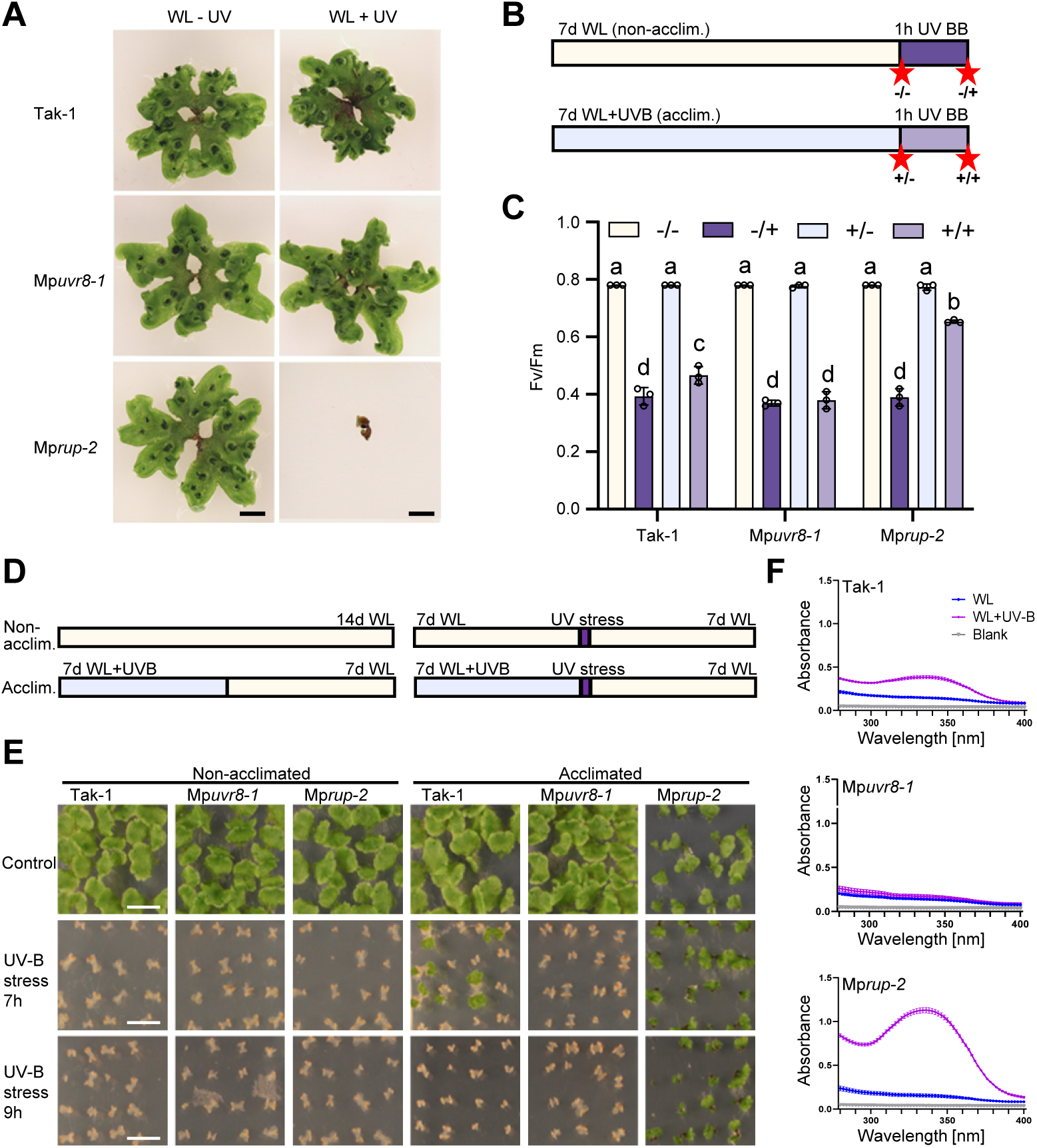
UV-B acclimation and survival is impaired in Mp*uvr8* and enhanced in Mp*rup*. **A)** Growth phenotype of 4-week-old wild-type (Tak-1), Mp*uvr8-1*, and Mp*rup-2* grown in white light supplemented (WL+UV) or not (WL-UV) with UV-B (1.5 μmol m^-2^ s^-1^). Bars = 1 cm. **B)** Scheme for UV-B treatment and sampling for F_v_/F_m_ measurements. Light yellow bar: white light (WL, 50 µmol m^-2^ s^-1^), non-acclimated condition; light blue bar: white light supplemented with narrowband UV-B (WL+UVB, 1.5 μmol m^-2^ s^-1^), acclimated condition; light/dark purple bars: broadband UV stress (UV BB, 25 μmol m^-2^ s^-1^). Red stars indicate timepoints of measurements: -/-, non-acclimated, non-stressed; -/+, non-acclimated, stressed; +/-, acclimated, non-stressed; +/+, acclimated, stressed. **C)** F_v_/F_m_ in 7-day-old wild-type (Tak-1), Mp*uvr8-1*, and Mp*rup-2* gemmalings grown and sampled as indicated in (B). Individual data points of biological replicates and means ± SD are shown (*n* = 3). Different letters indicate statistically significant differences between the means (P < 0.01) as determined by two-way ANOVA followed by Tukey’s post hoc honestly significant difference (HSD) test. **D)** Scheme for UV-B stress survival experiments. Plants were grown in WL or WL+UVB for 7 days, then exposed to 7 or 9 h of broadband UV-B stress, followed by recovery for 7 days in WL, after which they were imaged. **E)** UV-B stress survival in Tak-1, Mp*uvr8-1*, and Mp*rup-2* gemmalings grown as indicated in (D). Representative images are shown; bars = 1 cm. **F)** Absorbance spectra (280–400 nm) of methanolic extracts from 9-day-old Tak-1, Mp*uvr8-1*, and Mp*rup-2* gemmalings grown for 7 days in white light and exposed (WL+UV-B, purple lines) or not (WL, blue lines) to supplemental UV-B (1.5 μmol m^-2^ s^-1^) for 2 days. The gray line shows the absorbance of the buffer (blank). Values of independent measurements and means ± SEM are shown (*n* = 3).

To directly assess the effect of MpUVR8-mediated acclimation on photoprotection, we measured F_v_/F_m_ representing the maximum quantum yield of photosystem II (PSII) (Murchie and Lawson, 2013). Tak-1, Mp*uvr8*, and Mp*rup* plants were grown for 7 days under white light supplemented with narrowband UV-B (acclimated; +/-) or not (non-acclimated; -/-) (Fig. 3B). Acclimated and non-acclimated plants were then exposed to 1 h of broadband UV stress (+/+ and -/+, respectively). We measured F_v_/F_m_ before and after UV stress in both acclimated and non-acclimated plants (Fig. 3B). Non-stressed Tak-1, Mp*uvr8*, and Mp*rup* displayed an F_v_/F_m_ value close to 0.8 (Fig. 3C), which is characteristic of healthy Marchantia plants (Messant et al., 2023). When exposed to UV stress, F_v_/F_m_ decreased significantly and to the same extent in non-acclimated plants of all genotypes (Fig. 3C), suggesting photoinhibition resulting from PSII damage. However, applying the same UV stress treatment on acclimated plants revealed a UVR8-dependent photoprotective effect: in Tak-1 and even more so in Mp*rup*, F_v_/F_m_ following UV-B stress was higher in acclimated compared to non-acclimated plants, but not in Mp*uvr8* (Fig. 3C). Mp*UVR8.1^OE^*and Mp*UVR8.2^OE^* overexpression lines were similar to Mp*rup*, whereas Mp*RUP^OE^* overexpression lines phenocopied Mp*uvr8* (fig. S4B).

Next, we tested the survival rates of Tak-1, Mp*uvr8*, and Mp*rup* under UV-B stress. Plants were acclimated or not under narrowband UV-B for 7 days, followed by a 7- or 9-h broadband UV-B stress treatment, after which all plants were returned to white light for 7 days of recovery (Fig. 3D). Without acclimation, Tak-1, Mp*uvr8*, and Mp*rup* did not survive subsequent UV-B stress (Fig. 3E). Acclimated wild-type plants did not survive the 9-h UV-B stress treatment but some gemmalings resumed growth after 7 h of UV-B stress. Mp*rup* mutants recovered more than Tak-1 under both stress treatments (Fig. 3E and fig. S4C, D), indicating enhanced UV-B protection. By contrast, the Mp*uvr8* mutant did not survive the UV-B stress treatment even with prior UV-B acclimation, demonstrating that UV-B photoprotection was dependent on MpUVR8 (Fig. 3E). Complementing the Mp*uvr8-1* mutant with a *Pro*_Mp*UVR8*_:Mp*UVR8* construct restored UV-B stress survival in acclimated plants (fig. S4C). By contrast, overexpression of MpRUP (Mp*RUP^OE^*) phenocopied the Mp*uvr8* mutant phenotype (fig. S4D).

We prepared methanolic extracts from gemmalings and measured the absorption spectra between 280–400 nm to detect UV-absorbing compounds. There was a broad peak in UV-absorbance at ∼340 nm for UV-B-exposed Tak-1 plants, which was absent for the Mp*uvr8* mutant (Fig. 3F), indicating Marchantia plants accumulate UV-absorbing compounds in a UV-B- and UVR8-dependent manner. Consistent with their enhanced UV-B acclimatory response and resulting UV-B tolerance, extracts from UV-B-exposed Mp*rup* mutants had an even greater peak in UV absorption (Fig. 3F). Notably, the Mp*uvr8-1* Mp*rup-2* double mutant exhibited photoprotection and UV-absorbing capacity comparable to the Mp*uvr8-1* single mutant (fig. S5), demonstrating that the enhanced photoprotection and UV-B-absorbing capacity of Mp*rup* is dependent on MpUVR8. Next, we performed untargeted metabolomics of 9-day-old Tak-1, Mp*uvr8*, and Mp*rup* grown in white light for 7 days and exposed or not to supplemental UV-B for 2 days, following which we detected a total of 554 metabolites (table S1). Fifty-eight metabolites accumulated in response to UV-B in Tak-1 that were not observed in Mp*uvr8* (fig. S6A, table S2). Over half of these (30) were assigned to compounds from the shikimate-phenylpropanoid biosynthetic pathway, whereas metabolites from that pathway represented only about a quarter of all compounds detected (151/609) (fig. S6B and table S2). Unlike in Arabidopsis whereby sinapate esters play a major role in UV-B photoprotection (Leonardelli et al., 2024), no sinapate esters were detected in Marchantia (fig. S6C, table S1). However, UV-B-absorbing flavone glycosides (e.g. apigenin-7-glucoside, luteolin 7-O-glucuronide) accumulated in a MpUVR8-dependent manner under UV-B, and even more so in Mp*rup* (fig. S6C-E, table S2), consistent with previous reports (Clayton et al., 2018; Soriano et al., 2019). In addition to 28 of the 58 compounds that accumulated in response to UVR8 activation in Tak-1, 60 compounds did so specifically in Mp*rup* likely due to its hyperresponsiveness to UV-B (fig. S6A).

### Mp*cop1* mutants exhibit constitutive photomorphogenesis and UV-B tolerance

Interaction of UVR8 with the COP1/SPA E3 ligase complex and its subsequent inactivation is a key step in UV-B signal transduction in Arabidopsis (Favory et al., 2009; Yin et al., 2015; Lau et al., 2019). MpCOP1 and MpSPA interacted in yeast two-hybrid assays, both in the absence and presence of UV-B (Fig. 4A) (Zhang et al., 2021). By contrast, MpUVR8 interacted with MpCOP1 specifically under UV-B in yeast two-hybrid (Fig. 4A), LCI (Fig. 4B), and AP-MS assays (fig. S2).

**Figure 4.**
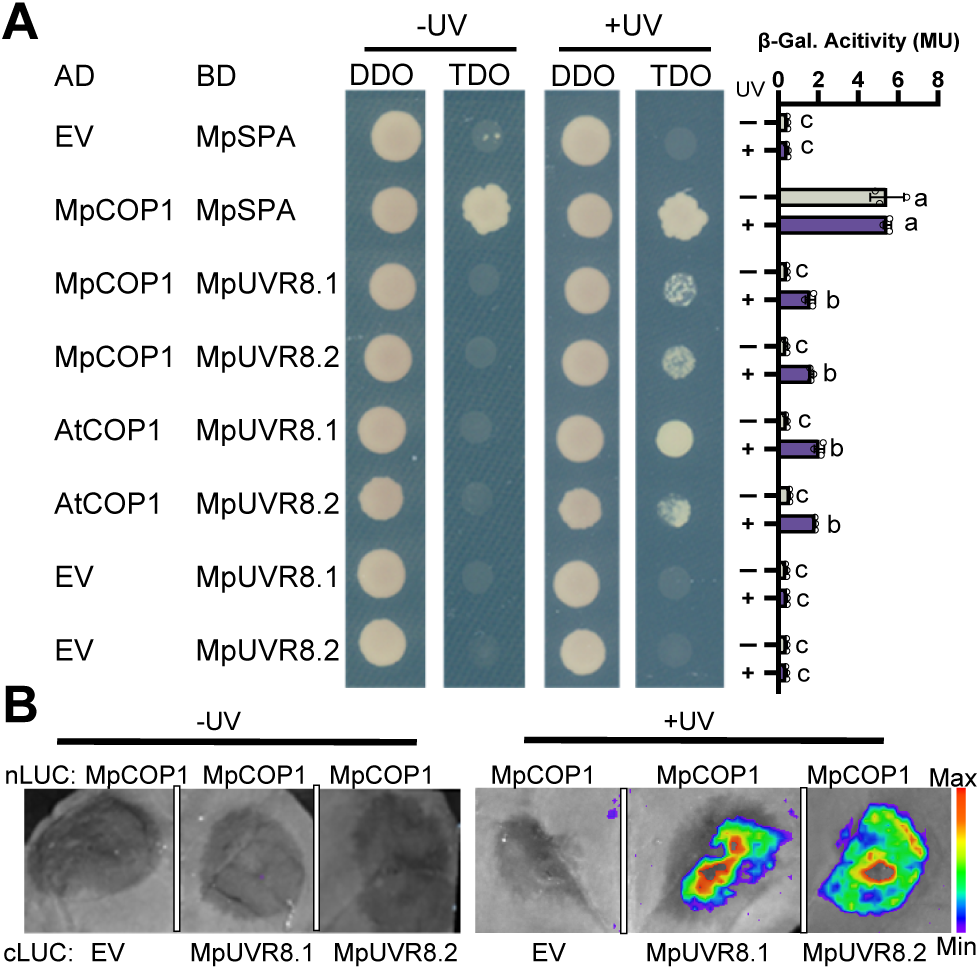
MpUVR8 and MpCOP1 interact in a UV-B-dependent manner. **A)** Yeast two-hybrid assay of the interactions of MpCOP1 with MpSPA and MpUVR8, as well as AtCOP1 with MpUVR8, in the presence (+UV) or absence (-UV) of UV-B. Left, growth assay; right, quantitative assay. Yeasts were grown on non-selective SD/-Trp/-Leu (DDO) or selective SD /-Trp/-Leu/-His (TDO) media. Individual data points of biological replicates and means ± SD are shown (*n* = 3). Different letters indicate statistically significant differences between the means (P < 0.01) as determined by one-way ANOVA followed by Tukey’s post hoc honestly significant difference (HSD) test. AD, Gal4 activation domain; BD, LexA DNA-binding domain; β-Gal., β-galactosidase; MU, Miller units; EV, empty vector. **B)** LCI assays of MpCOP1 with MpUVR8.1 and MpUVR8.2. N-terminal (nLUC) or C-terminal (cLUC) luciferase fusion proteins were expressed in *N. benthamiana* leaves and imaged 2 days after infiltration, with (+UV) or without (-UV) exposure to 20 min of broadband UV-B. An image of a representative leaf is shown.

Arabidopsis *cop1* null mutants have a seedling-lethal phenotype (McNellis et al., 1994) and no Marchantia Mp*cop1* mutant has been recovered until now (Zhang et al., 2021). We generated two viable Mp*cop1* null alleles using CRISPR/Cas9 mutagenesis (fig. S3). These mutants displayed stunted growth, delayed development, and reduced gemmae production compared to Tak-1 when grown under standard conditions (Fig. 5A). Whereas the Mp*cop1* mutants developed gemma cups and gemmae and thus could complete their asexual life cycle, they failed to complete their sexual life cycle because antheridiophores did not develop when induced under continuous white light supplemented with far-red light.

**Figure 5.**
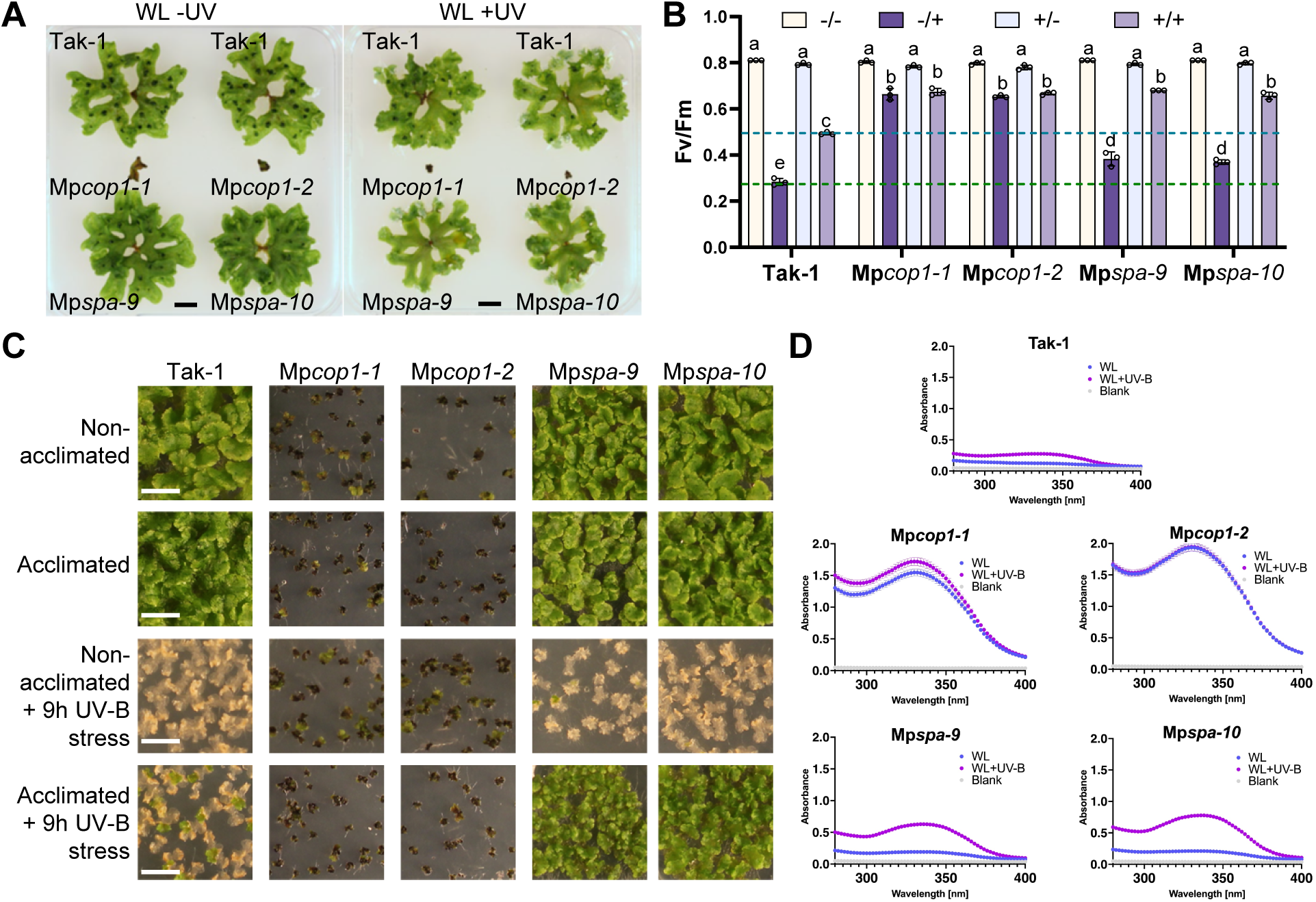
Mp*cop1* shows constitutive photomorphogenesis and UV-B tolerance, whereas Mp*spa* shows enhanced UV-B-induced UV-B tolerance. **A)** Growth phenotype of 4-week-old wild-type (Tak-1), Mp*cop1-1*, Mp*cop1-2*, Mp*spa-9*, and Mp*spa-10* plants grown in white light supplemented (WL +UV) or not (WL -UV) with UV-B. Scale bars = 1 cm. **B)** F_v_/F_m_ measurements in 7-day-old wild-type (Tak-1), Mp*cop1-1*, *Mpcop1-2*, Mp*spa-9*, and Mp*spa-10* gemmalings grown in white light (non-acclimated) or white light supplemented with UV-B (acclimated), without (non-stressed) or with (stressed) additional exposure to broadband UV-B stress for 60 min. -/-, non-acclimated, non-stressed; -/+, non-acclimated, stressed; +/-, acclimated, non-stressed; +/+, acclimated, stressed. Individual data points of biological replicates and means ± SD are shown (*n* = 3). Different letters indicate statistically significant differences between the means (P < 0.01) as determined by two-way ANOVA followed by Tukey’s post hoc honestly significant difference (HSD) test. **C)** UV-B stress survival in wild-type (Tak-1), Mp*cop1-1*, Mp*cop1-2*, Mp*spa-9*, and Mp*spa-10* plants grown for 7 days in white light supplemented (acclimated) or not (non-acclimated) with UV-B, then exposed or not to 9 h of broadband UV-B stress. Pictures were taken after a further 7-day recovery period in white light. **D)** Absorbance spectra (280–400 nm) of methanolic extracts from 9-day-old wild-type (Tak-1), Mp*cop1-1,* Mp*cop1-2*, Mp*spa-9*, and Mp*spa-10* gemmalings grown for 7 days in white light supplemented (WL+UV-B, purple lines) or not (WL, blue lines) with UV-B. The gray line shows the absorbance of the buffer (blank). Values of independent measurements and means ± SEM are shown (*n* = 3).

Mp*cop1* mutants exhibited constitutively enhanced photoprotection, as evidenced by their elevated F_v_/F_m_ ratios after UV-B stress (Fig. 5B). The strong constitutively photomorphogenic phenotype and enhanced photoprotection of Mp*cop1* agreed with how Mp*cop1* survived prolonged broadband UV-B stress, both with or without acclimation (Fig. 5C; note that the plants are viable as determined by non-stressed levels of F_v_/F_m_ after recovery). Absorption spectra of methanolic extracts also revealed constitutive hyperaccumulation of UV-absorbing compounds in Mp*cop1* (Fig. 5D). Thus, MpCOP1 activity as key repressor of photomorphogenesis is evident in Marchantia.

### Mp*spa* mutants show only a minor constitutively photomorphogenic growth phenotype, but strongly enhanced UV-B tolerance

To investigate the role of the single MpSPA protein, we generated two independent CRISPR/Cas9 knockout alleles (fig. S3). Surprisingly, both Mp*spa* alleles exhibited a phenotype comparable to Tak-1 under both white light and white light supplemented with UV-B (Fig. 5A). This is in stark contrast to the strong constitutively photomorphogenic phenotype observed for the Mp*cop1* mutant (Fig. 5A) and Arabidopsis *spa* quadruple mutants (Ordonez-Herrera et al., 2015).

Furthermore, we tested the UV-B acclimation response of Mp*spa* mutants. Mp*spa* showed enhanced basal UV-B tolerance when compared to wild type (“-/+”, Fig. 5B), as well as after UV-B acclimation (“+/+”, Fig. 5B). Mp*spa* mutants also showed an increased survival rate after prolonged UV-B stress if plants were previously acclimated to low UV-B (Fig. 5C). In accordance with their improved UV-B stress tolerance, Mp*spa* mutants constitutively accumulated more UV-absorbing compounds which became further pronounced upon UV-B-mediated acclimation (Fig. 5D).

As enhanced UV-B-acclimation of Mp*spa* resembled Mp*rup* mutants, we investigated whether MpSPA function was also associated with MpUVR8 redimerization. However, in contrast to Mp*rup*, Mp*spa* showed efficient MpUVR8 redimerization (fig. S7A). Additionally, overexpression of Mp*SPA* and Mp*COP1* reduced basal UV-B tolerance but did not abolish UV-B-induced photoprotection, in contrast to Mp*RUP* overexpression (fig. S7B-D). We conclude that MpCOP1 activity as a repressor of photomorphogenesis is not fully dependent on MpSPA, and that the absence of MpSPA enhances UV-B tolerance.

### MpUVR8 signaling components mediate transcriptional changes in response to UV-B

We analyzed the UVR8-mediated transcriptome in 7-day-old Marchantia gemmalings in response to 3 h of narrowband UV-B (tables S3 and S4). In Tak-1, we identified 480 and 114 genes up- and downregulated by UV-B, respectively (│log_2_ fold-change│ > 1; p-adj < 0.05), which all exhibited MpUVR8- and MpCOP1-dependent expression changes (tables S5A, B, and S6), thus representing the transcriptional MpUVR8 signaling output.

In Tak-1, Mp*rup*, and Mp*spa*, we identified a combined total of 756 UV-B-induced genes and 298 UV-B-repressed genes (│log_2_ fold-change│ > 1; p-adj < 0.05), with substantial overlaps between the genotypes (Fig. 6A-C, and tables S5A, B, and S6). Interestingly, induction or repression of transcripts after 3 h of UV-B was enhanced in Mp*rup* and Mp*spa* compared to Tak-1 (Fig. 6C), consistent with their UV-B hyper-responsive phenotypes (Fig. 3 and Fig. 5). Consistent with this, over two-thirds of genes that were up- or downregulated (│log_2_ fold-change│ > 1) specifically in Mp*rup* and Mp*spa* (“unique Mp*rup*” and “unique Mp*spa*”) were up- or downregulated in Tak-1 as well when considering a lower fold-change threshold (│log_2_ fold- change│ > 0 but <1; p-adj < 0.05) (Fig. 6A, B). Notably, whereas the transcriptomic profile of the Mp*cop1-1* mutant was unresponsive to UV-B, it resembled the profile of UV-B-responsive genotypes after UV-B exposure, with a large portion of genes constitutively up- or downregulated (Fig. 6C, D). These data support the role of MpCOP1 as a repressor of UV-B photomorphogenesis and COP1-inactivation by UVR8 as a key mechanism for UV-B-responsive gene expression changes.

**Figure 6.**
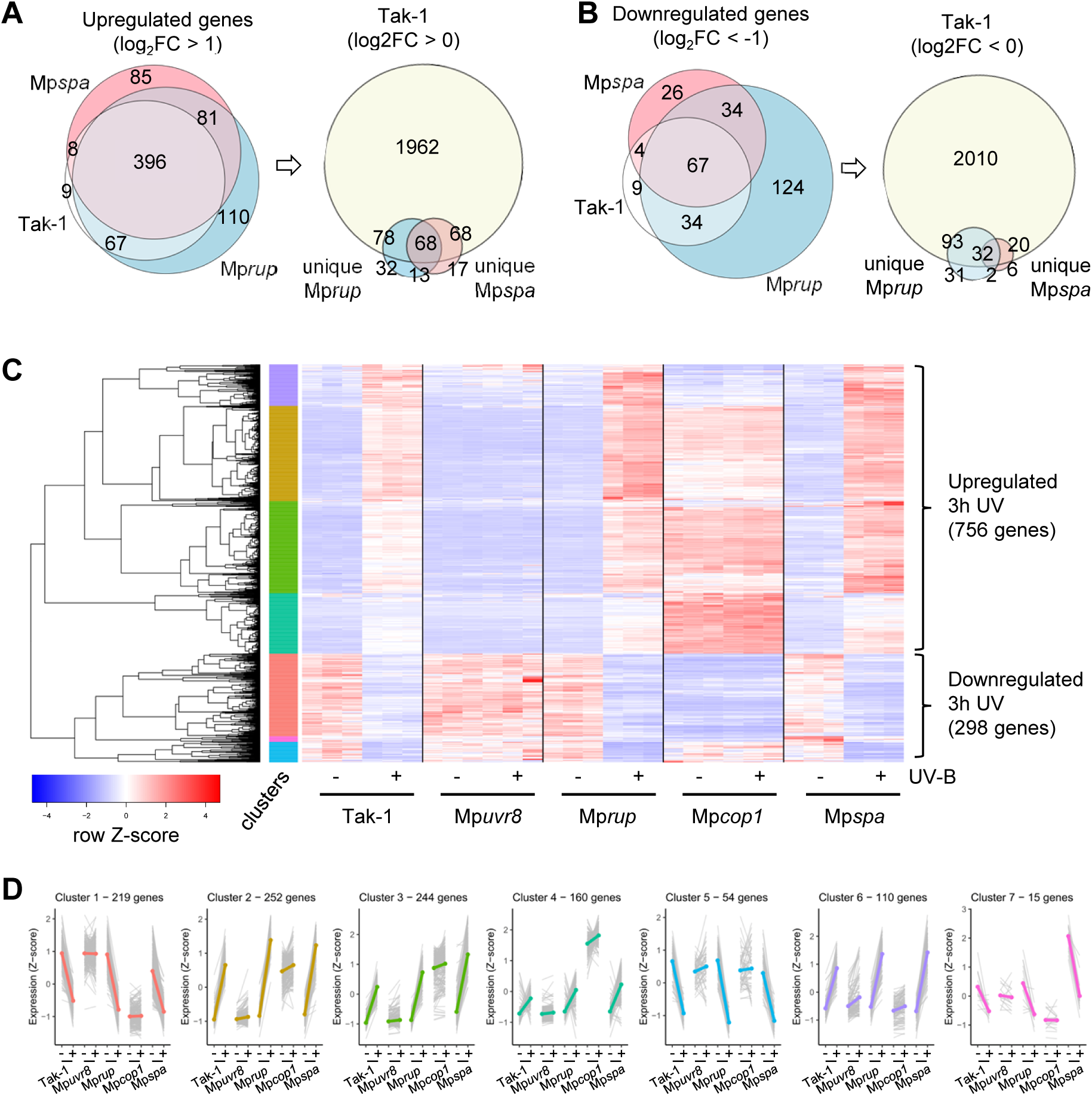
Transcriptome analysis reveals MpUVR8- and MpCOP1-dependent, UV-B-responsive gene regulation that is repressed by MpRUP and MpSPA. **A)** Euler plots showing overlaps between upregulated differentially expressed genes (DEGs) (p-adj < 0.05) upon exposure to 3 h UV-B in 7-day-old wild-type (Tak-1), Mp*rup*, and Mp*spa* gemmalings. Left: overlap between genes upregulated at least two-fold (log_2_FC > 1) in either Tak-1, Mp*rup*, or Mp*spa*. Right: overlap between upregulated (log_2_FC > 1) genes that are unique to Mp*rup* and/or Mp*spa*, and Tak-1 upregulated genes without a fold-change threshold (log_2_FC > 0). Corresponding gene lists: table S3a. **B)** Euler plots showing overlaps between downregulated DEGs (p-adj < 0.05) upon exposure to 3 h UV-B in 7-day-old Tak-1, Mp*rup*, and Mp*spa* gemmalings. Left: overlap between genes downregulated at least two-fold (log_2_FC < -1) in either Tak-1, Mp*rup*, or Mp*spa*. Right: overlap between downregulated (log_2_FC < -1) genes that are unique to Mp*rup* and/or Mp*spa*, and Tak-1 downregulated genes without a fold-change threshold (log_2_FC < 0). Corresponding gene lists: table S3b. **C)** Hierarchical clustering and heatmap of UV-B-regulated gene expression in Tak-1, Mp*uvr8-1,* Mp*rup-2,* Mp*cop1-1*, and Mp*spa-9* plants grown for 7 days under white light and exposed or not to supplemental UV-B for 3 h. All genes with a greater than two-fold change in expression in response to UV-B and a p-adj < 0.05 in at least one genotype are shown (total = 1005 genes). Red and blue colors indicate normalized read count Z-scores. The color bar on the left side indicates 7 major profiles (clusters) of gene expression. **D)** Gene expression profiles in clusters identified by hierarchical clustering. Z-scores for normalized read count values for each gene (gray lines) and for the cluster average (colored line) are shown.

Using hierarchical clustering, we defined seven major clusters of gene expression changes upon UV-B exposure (Fig. 6C, D, and table S6). Clusters 1 and 5 contain UV-B-downregulated genes, with gene expression in Mp*cop1* that is constitutively down- or upregulated, respectively. Cluster 7 consists of UV-B-downregulated genes that are more highly expressed in Mp*spa*. Clusters 4 and 6 contain genes that are induced after 3 h of UV-B in wild type, Mp*rup*, and Mp*spa*, but not in Mp*uvr8* and Mp*cop1*, and their constitutive expression in Mp*cop1* is either elevated or comparable in comparison to that in wild type, respectively. Clusters 2 and 3 contain genes induced after 3 h of UV-B exposure in wild type, with hyper-induced expression in Mp*spa* and Mp*rup*, respectively. We profiled the expression of components of the UVR8 pathway in the data: Mp*HY5*, Mp*RUP*, and Mp*COP1* (cluster 2) were upregulated upon UV-B in wild type (table S6), whereas Mp*UVR8* was constitutively expressed.

### MpSPA plays a minor role in repressing visible light photomorphogenesis compared to MpCOP1

In Arabidopsis, *spa1234* (*spaQn*) quadruple mutants undergo constitutive photomorphogenesis with a phenotype similar to strong *cop1* mutants (such as *cop1-1*) and stronger than the weak *cop1-4* allele (Fig. 7A) (McNellis et al., 1994; Laubinger et al., 2004; Ordonez-Herrera et al., 2015). However, Marchantia Mp*spa* mutants exhibited only a minor growth phenotype in white light compared to the strongly affected Mp*cop1* mutants (Fig. 7A, Fig. 5A). Thus, we investigated the transcriptome profile Mp*spa* and Mp*cop1* mutants in more detail.

**Figure 7.**
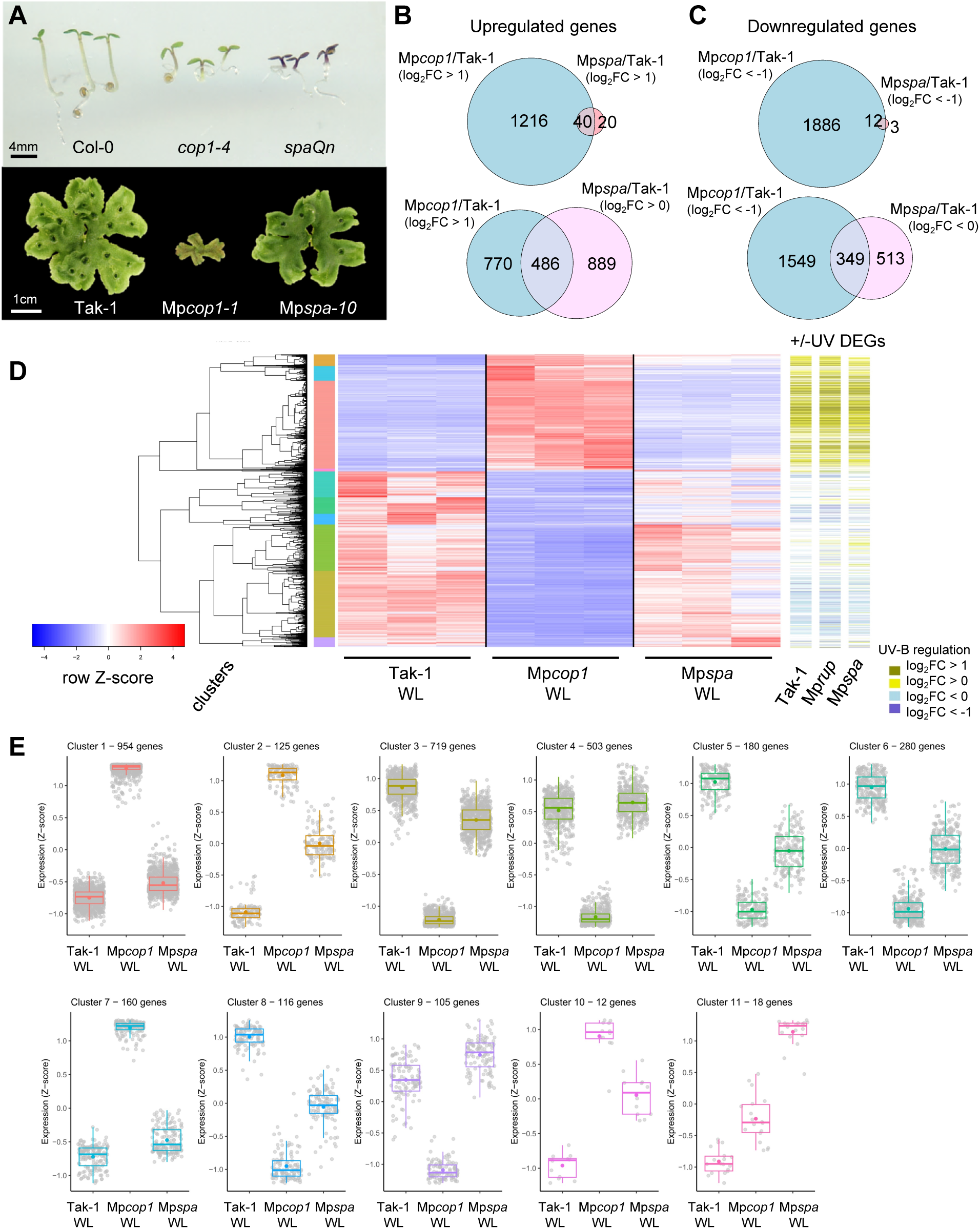
MpSPA plays a minor role in repressing Marchantia photomorphogenesis under visible light. **A)** Representative images of 4-day-old Arabidopsis seedlings of wild-type (Col-0), *cop1-4*, and *spaQn* mutants, and 28-day-old Marchantia thalli of wild-type (Tak-1), Mp*cop1-1*, and Mp*spa-10* grown in white light. Scale bars: 4 mm or 1 cm, as indicated. **B)** Euler plots showing overlaps between upregulated DEGs (p-adj < 0.05) in Mp*cop1* and Mp*spa* mutants compared to Tak-1 wild type. Gemmalings were grown for 7 days in white light. Top: overlap between genes upregulated at least two-fold (log_2_FC > 1) in either Mp*cop1*/Tak-1 or Mp*spa*/Tak-1. Bottom: overlap between genes upregulated at least two-fold (log_2_FC > 1) in Mp*cop1*/Tak-1 and genes upregulated without a fold-change threshold (log_2_FC > 0) in Mp*spa*/Tak-1. Corresponding gene lists: table S3c. **C)** Euler plots showing overlaps between downregulated DEGs (p-adj < 0.05) in Mp*cop1* and Mp*spa* mutants compared to Tak-1 wild type. Gemmalings were grown for 7 days in white light. Top: overlap between genes downregulated at least two-fold (log_2_FC < -1) in either Mp*cop1*/Tak-1 or Mp*spa*/Tak-1. Bottom: overlap between genes downregulated at least two-fold (log_2_FC < -1) in Mp*cop1*/Tak-1 and genes downregulated without a fold-change threshold (log_2_FC < 0) in Mp*spa*/Tak-1. Corresponding gene lists: table S3d. **D)** Hierarchical clustering and heatmap of gene expression in Tak-1, Mp*cop1-1*, and Mp*spa-9* plants grown for 7 days in white light. All genes with a greater than two-fold change in expression between mutants and wild type and a p-adj < 0.05 in at least one comparison are shown (total = 2975 genes). Red and blue colors indicate normalized read count Z-scores. The color bar on the left side indicates 11 major profiles (clusters) of gene expression. The color bars on the right indicate if genes are significantly regulated after UV-B exposure in Tak-1, Mp*rup*, and Mp*spa*. **E)** Gene expression profiles in clusters identified by hierarchical clustering. Z-scores for normalized read count values for each gene are shown as gray points; their distribution is represented as a colored box plot.

By comparing white light–grown Mp*cop1* and Mp*spa* versus Tak-1, we found that in Mp*cop1*, 1,256 and 1,898 genes were significantly up- or downregulated (│log_2_ fold-change│ > 1; p-adj < 0.05), respectively (Fig. 7B, C). By stark contrast, only 60 genes were upregulated and 15 genes were downregulated in Mp*spa* using the same threshold criteria (Fig. 7B, C, and table S5C, D), which is significantly different from Arabidopsis, where over 7,000 differentially expressed genes (DEGs) were observed between the *spaQ* mutant and the wild type (Pham et al., 2020). Of those genes dysregulated in Mp*spa*, 40 out of 60 and 12 out of 15 genes were also up- and downregulated, respectively, in Mp*cop1*. We further checked if genes dysregulated in Mp*cop1* but not in Mp*spa* were in fact dysregulated in Mp*spa* but at a lower fold-change threshold. Indeed, out of the 1,256 upregulated genes in Mp*cop1*, 486 were shared with Mp*spa* when considering a looser threshold for genes upregulated in Mp*spa* (│log_2_ fold-change│ > 0; p-adj < 0.05; Fig. 7B). Similarly, out of the 1,898 downregulated genes in Mp*cop1*, 349 were also downregulated in Mp*spa* (Fig. 7C). This indicates a larger overlap between genes dysregulated in Mp*cop1* and Mp*spa* mutants in comparison to wild type, but also identifies clear differences between MpCOP1- and MpSPA-dependent genes at the genome-wide level.

A broader examination of gene expression levels across wild type, Mp*cop1*, and Mp*spa* genotypes confirmed that, although the overall profile of Mp*spa* was similar to wild-type, many transcripts had levels that were intermediate between Tak-1 and Mp*cop1* (Fig. 7D and table S7). Using hierarchical clustering, we defined 11 main clusters of gene expression (Fig. 7E), specifically four for Mp*cop1*-upregulated genes (clusters 1, 2, 7 and 10), six for Mp*cop1*-downregulated genes (clusters 3, 4, 5, 6, 8 and 9), and a small cluster of genes specifically upregulated in Mp*spa* (cluster 11). Interestingly, the cluster groups defined by Mp*cop1*-up- and downregulated patterns of gene expression could be discriminated by the extent to which expression in Mp*spa* aligned with the expression in Mp*cop1* (Fig. 7E).

Finally, since we observed that the regulation of many genes was dependent on MpCOP1 under white light, and we established that MpCOP1 has a conserved role as a repressor of photomorphogenesis, we tested if genes dysregulated in Mp*cop1* were induced by UV-B in UV- B-responsive genotypes. We observed that most genes upregulated under white light in Mp*cop1* were induced by UV-B in Tak-1, Mp*rup*, or Mp*spa* (Fig. 7D), whereas many genes constitutively downregulated in Mp*cop1* were repressed by UV-B. However, some of the Mp*cop1*-downregulated genes were induced after 3 hours of UV-B, suggesting more complex dynamics between induction under UV-B and steady-state levels in Mp*cop1*.

Overall, our analysis shows that whereas MpCOP1 plays a major role in repressing the light-inducible transcriptome of Marchantia, MpSPA plays a minor yet detectable role.

### MpHY5 promotes MpUVR8 signaling

The induction of Mp*HY5* expression is UV-B- and MpUVR8-dependent, repressed by both MpRUP and MpSPA, and Mp*HY5* is constitutively upregulated in the Mp*cop1-1* mutant (table S6). To test the role of MpHY5 in Marchantia, we generated Mp*hy5* knockout mutants by CRISPR/Cas9 (fig. S3). We further investigated the photoprotection and UV-B-absorbing capacity of Mp*hy5* mutants. Mp*hy5* mutants showed slightly reduced basal photoprotection, as determined by measuring the F_v_/F_m_ ratio after UV-B stress exposure (-/+; Fig. 8A). Interestingly, and in contrast to Arabidopsis *hy5 hyh* double mutants (Leonardelli et al., 2024), Mp*hy5* mutants were able to acclimate to UV-B (+/+ versus -/+; Fig. 8A), but Mp*hy5* mutants nevertheless showed a significantly reduced capacity to protect their photosynthetic machinery from subsequent UV-B-induced damage compared to UV-B-acclimated wild type (+/+; Fig. 8A). In agreement, UV-B-acclimated Mp*hy5* mutants were less tolerant to 7 h of UV-B stress than Tak-1 (Fig. 8B). We further measured the UV-B absorbance profiles of methanolic extracts from different genotypes and found that, consistent with the reduced photoprotection observed in UV-B-exposed Mp*hy5*, Mp*hy5* accumulated slightly less UV-B-absorbing compounds compared to Tak-1 (Fig. 8C).

**Figure 8.**
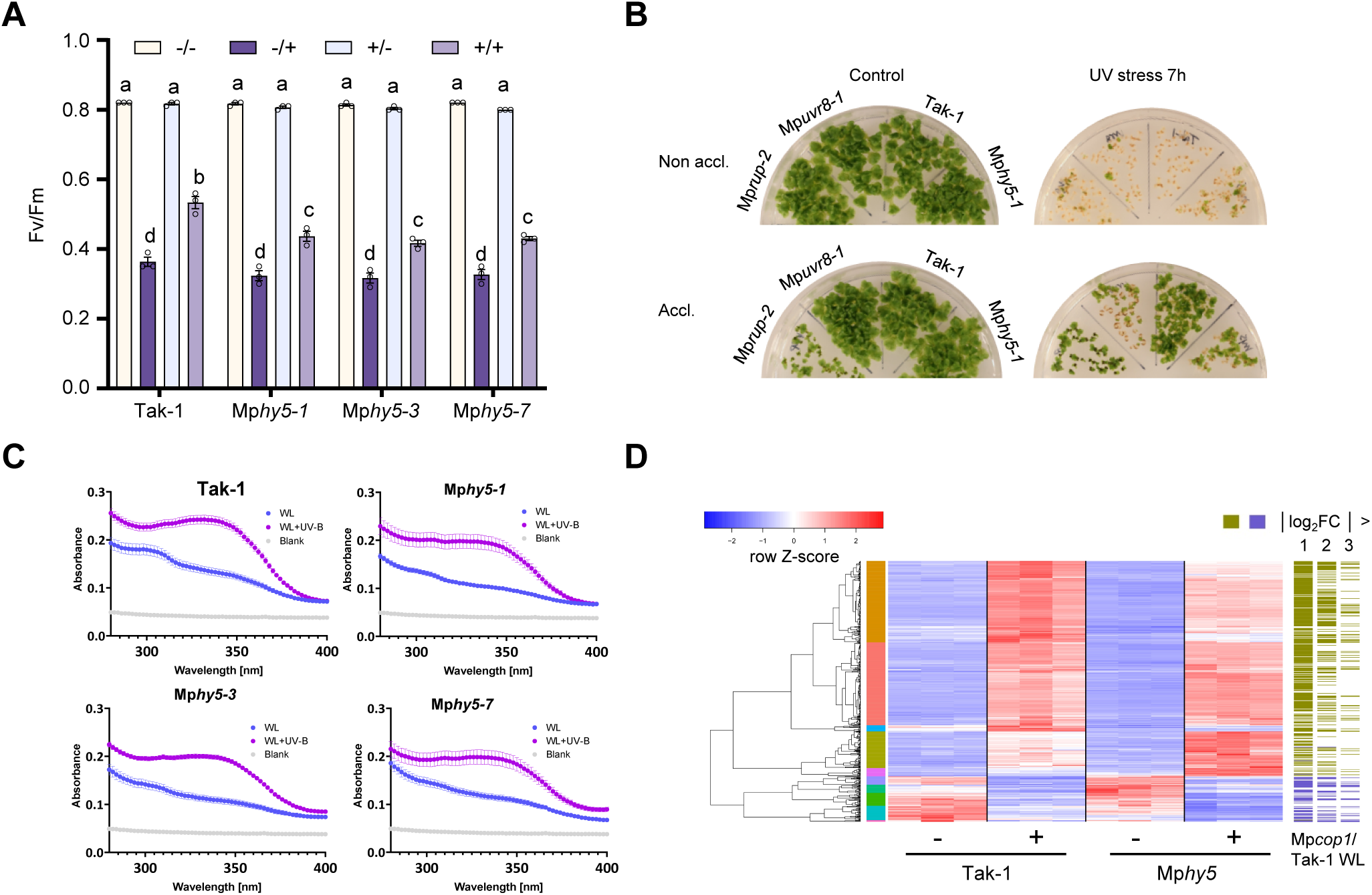
MpHY5 is involved in UV-B and visible light photomorphogenesis. **A)** F_v_/F_m_ measurements in wild-type (Tak-1), Mp*hy5-1*, Mp*hy5-3*, and Mp*hy5-7* plants grown for 7 days in white light (non-acclimated) or white light supplemented with UV-B (acclimated), without (not stressed) or with (stressed) additional exposure to broadband UV-B stress for 60 min. -/-, not acclimated, not stressed; -/+, not acclimated, stressed; +/-, acclimated, not stressed; +/+, acclimated, stressed. Individual data points of biological replicates and means ± SD are shown (*n* = 3). Letters indicate statistically significant differences between the means (P < 0.01) as determined by two-way ANOVA followed by Tukey’s post hoc honestly significant difference (HSD) test. **B)** UV-B stress survival of Tak-1, Mp*hy5-1*, Mp*rup-2*, and Mp*uvr8-1* plants grown for 7 days in white light supplemented (Accl.) or not (Non-accl.) with UV-B, then exposed to 7 h of broadband UV-B stress (UV stress 7h) or not (Control). Pictures were taken after recovery for another 7 days in white light. **C)** Absorbance spectra (280–400 nm) of methanolic extracts from 9-day-old Tak-1, Mp*hy5-1*, Mp*hy5-3*, and Mp*hy5-7* gemmalings grown for 7 days in white light and exposed (WL+UV-B, purple lines) or not (WL, blue lines) to supplemental UV-B (1.5 μmol m^-2^ s^-1^) for 2 days. The gray lines show absorbance of the buffer (blank). Values of independent measurements and means ± SEM are shown (*n* = 3). **D)** Hierarchical clustering and heatmap of UV-B-regulated gene expression in Tak-1 and Mp*hy5-1* plants grown for 7 days and exposed, or not, to supplemental UV-B for 3h. All genes with a greater than two-fold change in expression under UV-B and a p-adj < 0.05 in at least one genotype are shown (total = 501 genes). Red and blue colors indicate normalized read count Z-scores. The color bar on the left side indicates 10 major profiles (clusters) of gene expression. The color bars on the right indicate if genes are significantly misregulated in white light in Mp*cop1* compared to Tak-1, at varying fold-change thresholds.

To further understand how MpHY5 promotes MpUVR8 signaling, we performed another RNA-seq experiment with 7-day-old Tak-1 and Mp*hy5-1* Marchantia gemmalings exposed or not to 3 h of narrowband UV-B (tables S3, S4, S8, and S9). Using heatmap analysis and hierarchical clustering, we examined the 501 genes regulated upon UV-B exposure in Tak-1 and/or Mp*hy5-1* (│log2 fold-change│ > 1; p-adj < 0.05) (table S9). This analysis revealed 10 major clusters of gene expression changes in response to UV-B exposure (Fig. 8D, and fig. S8). Notably, few genes were entirely dependent on Mp*HY5*, though certain trends were observed where some genes showed reduced or enhanced UV-B-induced expression compared to Tak-1. Mp*HY5-1* (one base deletion, see fig. S3) and Mp*COP1* were hyperinduced in Mp*hy5-1* (cluster 3), while the induction of Mp*RUP* was unaffected by the Mp*hy5-1* mutation (cluster 1). Most UV-B-induced genes were constitutively upregulated in Mp*cop1-1* (Fig. 8D and table S5), and we noticed that the most MpHY5-dependent genes (clusters 2 and 7) were associated with the highest levels of constitutive upregulation in Mp*cop1*, while the genes that showed the most hyperinduction in Mp*hy5* were associated with the lowest constitutive expression levels in Mp*cop1* (Fig. 8D, fig. S9, and table S8). This is in agreement with an evolutionary conservation of the COP1-HY5 regulatory module in Marchantia photomorphogenesis. Taken together, our data indicate that MpHY5 acts antagonistically to MpCOP1 and is involved in mediating UVR8-responsive gene expression in Marchantia, although to a much lesser extent than that facilitated by HY5 and HYH in Arabidopsis, suggesting that other transcription factors play major roles in UVR8 signaling in Marchantia.

## DISCUSSION

UV-B exposure elicits UVR8-dependent transcriptional changes underlying UV-B acclimation phenotypes in plants (Kliebenstein et al., 2002; Brown et al., 2005; Favory et al., 2009; Tilbrook et al., 2016; Tavridou et al., 2020a; Hu et al., 2024; Dannay et al., 2025). The present study defines the MpUVR8-mediated genome-wide transcriptional response and provides physiological evidence for how MpUVR8 regulates acclimation and UV-stress tolerance in Marchantia using a broad panel of knock-out mutants in the UVR8 signaling pathway. Importantly, our study further demonstrates that although core components of UV-B signaling are conserved, their regulatory interactions and functions have significantly diversified.

In contrast to that in Arabidopsis and the moss *Physcomitrium patens* (hereafter Physcomitrium), MpUVR8 has been reported to form weaker dimers and increased levels of monomer *in vivo* in the absence of UV-B, which is observed when analyzing MpUVR8 dimer/monomer ratio by SDS-PAGE and immunoblotting using samples that are not heat-denatured before electrophoresis (Soriano et al., 2018). Although this type of assay is accurate for the dimer/monomer status of wild-type AtUVR8 (Rizzini et al., 2011; Heijde and Ulm, 2013; Heilmann and Jenkins, 2013; Findlay and Jenkins, 2016; Moriconi et al., 2018), it can make weak dimeric UVR8 mutant variants appear monomeric, whereas *in vitro* assays (Christie et al., 2012; Wu et al., 2012; Heijde et al., 2013) as well as more stringent assays using protein-protein crosslinking identify these variants as homodimeric *in vivo* (Rizzini et al., 2011; Podolec et al., 2021b). Thus, we employed crosslinking to show that MpUVR8 is indeed dimeric *in vivo* in the absence of UV-B, consistent with the UV-B-dependent dimer-to-monomer transition being a conserved feature of the UVR8 photocycle (Rizzini et al., 2011; Tilbrook et al., 2016; Podolec et al., 2021a). Furthermore, we provide evidence that negative feedback regulation through MpRUP-facilitated redimerization of MpUVR8 is conserved in Marchantia. Consistent with this, the Mp*rup* mutant displayed a strong UV-B photomorphogenic phenotype, characterized by growth inhibition resulting in a small size, a strong pigmentation of the thallus, and higher UV-B stress tolerance associated with increased accumulation of UV-absorbing compounds (see also Clayton et al., 2018). Interestingly, this conditional phenotype of Mp*rup* specifically under UV-B phenocopies the constitutive Mp*cop1* phenotype, in agreement with active UVR8-mediated inactivation of COP1 underlying UV-B signaling (Yin et al., 2015; Lau et al., 2019). We further show that GFP-MpUVR8 expressed under its own promoter localizes mainly to the cytosol and displays UV-B-dependent nuclear accumulation in Marchantia, confirming findings using a *Citrine-*Mp*UVR8* overexpression line (Kondou et al., 2019). Thus, we conclude that the homodimeric ground state of UVR8, its UV-B dependent monomerization, nuclear accumulation, and COP1 inactivation, as well as negative feedback regulation of UVR8 through RUP-facilitated redimerization are well-conserved in Marchantia.

In flowering plants, COP1 forms an E3 ligase complex together with SPA proteins, which together function as suppressors of photomorphogenesis; i.e. COP1 requires SPA proteins to function (Hoecker, 2017; Podolec and Ulm, 2018). Since non-plant COP1 orthologs are not regulated by light, it is hypothesized that the evolution of SPA proteins was a crucial step in enabling photoreceptor-mediated regulation of COP1/SPA complex activity (Ranjan et al., 2014; Balcerowicz et al., 2017; Podolec and Ulm, 2018). Active phytochrome and cryptochrome photoreceptors directly interact with SPA proteins, which represses the E3 ligase activity of the COP1/SPA complex (Hoecker, 2017; Podolec and Ulm, 2018). In agreement, Arabidopsis quadruple *spa* mutants exhibit strong constitutive photomorphogenesis (Ordonez-Herrera et al., 2015). However, it is of note that *spa* quadruple null mutants can complete their life cycle whereas *cop1* null mutants cannot (McNellis et al., 1994; Ordonez-Herrera et al., 2015), suggesting that COP1 can function partially in a SPA-independent manner. COP1, but not SPA proteins, are also necessary for transient gene expression changes upon exposure to UV-B (Favory et al., 2009). Recently, the role of COP1/SPA has started to be explored in more phylogenetically distant land plant lineages such as bryophytes, including Physcomitrium (Kreiss et al., 2023). A Physcomitrium Pp*cop1_9x* mutant lacking all nine PpCOP1(a-i) orthologs was viable and displayed a rather minor growth phenotype, certainly much less severe than the Arabidopsis *cop1* mutant phenotypes (Kreiss et al., 2023). In this study, we analyzed mutants of the single Mp*COP1* gene in light-grown Marchantia plants and found that, although they completed their asexual life cycle, they exhibited profound transcriptome changes and a severe constitutively photomorphogenic phenotype. Additionally, consistent with the described role of COP1 in UV-B signaling in Arabidopsis and Chlamydomonas (Oravecz et al., 2006; Favory et al., 2009; Allorent et al., 2016; Tilbrook et al., 2016), transient gene expression changes upon exposure to UV-B were dependent on MpCOP1. The Marchantia genome contains a single gene encoding MpSPA that interacts with MpCOP1 and MpCRY (Zhang et al., 2021). Surprisingly, Mp*spa* null mutants exhibited at most a mild constitutively photomorphogenic phenotype in white light, unlike Mp*cop1*. Interestingly, in Physcomitrium, a Pp*spa_ab* double mutant displayed an intermediate phenotype between that of the wild type and Pp*cop1_9x*. This suggests that SPA proteins indeed play a lesser role in repressing light signaling in bryophytes compared to in angiosperms (Huang et al., 2013; Ordonez-Herrera et al., 2015; Artz et al., 2019; Kreiss et al., 2023) and that high dependence of COP1 on the accessory SPA proteins evolved later, meaning that SPA protein evolution is separate to the evolution of COP1 activity regulation by light. This is further supported by the discovery of the UVR8- and cryptochrome photoreceptor–mediated COP1 inactivation mechanism of high-affinity competitive binding and displacement of COP1 substrates (Lau et al., 2019; Ponnu et al., 2019; Trimborn et al., 2025). Moreover, complementation of corresponding Arabidopsis mutant phenotypes was possible with a Physcomitrium COP1 ortholog but not a SPA ortholog (Ranjan et al., 2014), further indicating the divergence of SPA function between evolutionary lineages.

Enhanced UV-B-induced changes in gene expression, increased accumulation of UV-absorbing compounds, and enhanced UV-B acclimation and photoprotection against UV stress in Mp*spa* mutants compared to that in wild type revealed a repressor role for SPA in the UV-B signaling pathway of Marchantia. This is similar to Mp*rup* mutants, although long-term growth under UV-B was not as strongly inhibited in Mp*spa*. MpSPA also did not facilitate MpUVR8 redimerization like observed for MpRUP. Several hypotheses can be proposed to explain the Mp*spa* phenotype. It has been reported that MpSPA can promote the ubiquitination of MpHY5 by MpCOP1 (Zhang et al., 2021). Consequently, the lack of MpSPA could result in reduced ubiquitination of MpHY5 by MpCOP1, leading to higher activation of downstream genes. Also, UV-B-activated MpUVR8 may inactivate MpCOP1 more strongly in the absence of MpSPA, leading to an enhanced UV-B response. MpSPA might function by “shielding” the MpCOP1/SPA complex from being inhibited by MpUVR8 under UV-B by providing an additional UVR8-irrepressible activity. Moreover, it has been suggested that SPA proteins promote the phosphorylation of several proteins involved in light signaling in Arabidopsis, including UVR8 and HY5 (Wang et al., 2021; Liu et al., 2024). Therefore, in the event such activity is conserved, the absence of MpSPA may affect the phosphorylation status and activity of light signaling components, enabling the Mp*spa* mutant to respond more strongly to UV-B and to produce more “sunscreen” metabolites. Future work is required to elucidate the precise mechanism by which MpSPA exerts its repressive effect on UV-B signaling in Marchantia, and to determine if a similar SPA protein function exists in Arabidopsis or whether this represents a species-specific UV-B response regulatory mechanism. Overall, genetic analysis of COP1/SPA function suggests that, whereas the role of COP1 is widely conserved within the green lineage, the importance of SPA proteins in regulating COP1 activity and repressing photomorphogenesis increased during plant evolution.

It is of note that the role and importance of the transcriptional regulators HY5/HYH in UVR8-signaling has also changed during plant evolution. HY5 is a major player in UVR8-signaling in Arabidopsis (Ulm et al., 2004; Brown et al., 2005; Oravecz et al., 2006; Binkert et al., 2014), whereas the closest Chlamydomonas homolog of HY5, CrBLZ3, plays only a minor role in the algal UV-B response (Dannay et al., 2025). Instead, the B-box family transcription factor and CrCOP1 substrate CrCONSTANS is the main player in the CrUVR8 signaling pathway (Gabilly et al., 2019; Tokutsu et al., 2019; Dannay et al., 2025). In Marchantia, MpHY5 controls a subset of MpUVR8-induced genes but has a less prominent role than its counterpart in Arabidopsis. Other transcription factor(s) mediating MpUVR8 signaling remain to be identified.

In conclusion, our study provides a comprehensive understanding of the UV-B perception and signaling mechanisms in the liverwort model organism Marchantia. Our findings further confirm and strengthen the notion that the UVR8 photoreceptor and its associated signaling components are conserved in Marchantia, highlighting the evolutionary importance of this pathway in mediating UV-B responses across land plants. Our analysis of the COP1/SPA E3 ligase complex and its HY5 substrate in Marchantia also revealed divergences in the functions of this photomorphogenesis-repressing complex.

## MATERIALS AND METHODS

### Plant materials

The *Marchantia polymorpha* accessions Takaragaike-1 (Tak-1) and Tak-2 were used as male and female wild type, respectively (Ishizaki et al., 2008). The Mp*uvr8-1* mutant was described previously (Kondou et al., 2019), whereas the following mutants were generated using CRISPR– Cas9-mediated mutagenesis (Sugano et al., 2018; Sauret-Güeto et al., 2020): Mp*rup-2,* Mp*rup-3,* Mp*cop1-1,* Mp*cop1-2,* Mp*spa-9,* Mp*spa-10,* Mp*hy5-1,* Mp*hy5-3,* and Mp*hy5-7*. Two different gene-specific guide RNAs (gRNAs, see table S10) were selected using the CRISPRdirect (https://crispr.dbcls.jp/) or CasFinder (https://marchantia.info/tools/casfinder/) tools (Aach et al., 2014). Corresponding complementary DNA oligos were annealed and ligated into the BsaI sites of pMpGE_En03, then subcloned to the binary vector pMpGE010 (Sugano et al., 2018) using Gateway LR Clonase II (Thermo Fisher Scientific). To generate the Mp*hy5* mutants, complementary oligos were inserted into BbsI sites of the L1_lacZgRNA-Ck2 and L1_lacZgRNA-Ck3 vectors, respectively, after which these were recombined by SapI restriction enzymes with the L1_HyR-Ck1 and L1_Cas9-Ck4 vector into the pCsA acceptor plasmid using Golden Gate Loop Assembly (Sauret-Güeto et al., 2020). Following transformation, frame-shift and early stop codon mutants were identified using PCR genotyping and Sanger sequencing (table S10 and fig. S3).

To generate *Pro*_Mp*UVR8*_:*GFP*-Mp*UVR8* reporter lines, we modified the previously described pMpGWB301-MpUVR8 genomic construct (Kondou et al., 2019). The vector was digested with MluI and HpaI and the GFP coding sequence was ligated immediately upstream of the Mp*UVR8* start codon. To create Pro_Mp*CHS2*_:*LUC* reporter lines, the predicted Mp*CHS2* promoter including 5ʹUTR sequence (3,000 bp region upstream of the Mp*CHS2* start codon) was amplified from Tak-1 genomic DNA and the PCR product was inserted into pDONR207 via Gateway BP reaction. The *Pro*_Mp*CHS2*_ sequence was then transferred into the pMpGWB331 vector by LR reaction, followed by transformation into the Tak-1 wild-type background. To create *Pro*_Mp*CHS2*_*:LUC* lines in the Mp*uvr8-1* and Mp*rup-1* backgrounds, Tak-1/*Pro*_Mp*CHS2*_*:LUC* was first crossed with Tak-2 to obtain Tak-2/*Pro*_Mp*CHS2*_*:LUC* in female background, and then the male Mp*uvr8-1* or Mp*rup-2* mutants were crossed with the female Tak-2/*Pro*_Mp*CHS2*_*:LUC* line. To generate overexpression lines in Tak-1, the coding sequences including stop codons of Mp*UVR8.1*, Mp*UVR8.2*, Mp*RUP*, Mp*COP1*, and Mp*SPA* were first cloned into the pDONR207 entry vector via BP reaction, then transferred via LR reaction into the pMpGWB336 destination vector containing the *ELONGATION FACTOR 1α* (Mp*EF1α*, Mp3g23400) promoter and an N-terminal Red Fluorescent Protein (RFP) tag. Agrobacterium-mediated Marchantia thallus transformation was performed as previously described (Kubota et al., 2013). Marchantia spore transformation was used to generate Mp*hy5* mutants as previously described (Chiyoda et al., 2008).

Mp*uvr8-1* Mp*rup-2* Cas9-free double mutants were generated by crossing Mp*rup-2* (male) with Tak-2 to select for female lines, followed by crossing Mp*uvr8-1* (male) with Mp*rup-2* (female), as previously described (Mulvey and Dolan, 2023). For those plants where the inheritance of the parental alleles was confirmed, replica plating on plates with and without hygromycin was repeated to further verify the absence of the Cas9 transgene.

### Growth conditions and light treatments

For maintenance, Marchantia plants were cultured aseptically on half-strength Gamborg’s B5 medium with 1.2% (w/v) agar at 22°C under continuous white light (50 μmol m^-2^ s^-1^) in a CLF GroBank growth chamber. For UV-B acclimation experiments, continuous white light was supplemented with narrowband UV-B tubes (Philips TL20W/01RS, 1.5 μmol m^-2^ s^-1^). For UV-B stress treatments, plates were irradiated under broadband UV-B tubes (Philips TL20W/12RS; 25 μmol m^-2^ s^-1^).

### Crosslinking and immunoblotting

Crosslinking of proteins and immunoblot analysis of 7-day-old Marchantia gemmalings was performed following a modified version of a previously described method (Rizzini et al., 2011), except that the extraction buffer did not contain leupeptine, dichloroisocumarin, MG115, and PSI. Polyclonal rabbit anti-MpUVR8^(3-17)^ antibodies raised and purified against a synthetic peptide (amino acids C + 3–17: CDSKMSSNERPRRRVL) were used as the primary antibodies (Eurogentec), with horseradish peroxidase (HRP)-conjugated anti-rabbit immunoglobulins (Dako A/S) as secondary antibodies. The signal was revealed using ECL Select Western Blotting Detection Reagent (GE Healthcare) and analyzed using an Amersham Imager 680 camera system (GE Healthcare).

### Quantitative reverse transcription PCR (RT-qPCR) analysis

RNA was isolated with the RNeasy Plant Mini kit (Qiagen) according to the manufacturer’s instructions and was treated on-column with the RQ1 RNase-Free DNase (Promega), before serving as template for cDNA synthesis using the TaqMan Reverse Transcription Reagents kit (Thermo Fisher Scientific) with a 1:1 mix of oligo-dT and random hexamer primers. RT-qPCR was performed using the PowerUp SYBR Green Master Mix (Thermo Fisher Scientific) on a QuantStudio^TM^ 5 Real-Time PCR System (Thermo Fisher Scientific). Mp*EF1α* was used as reference gene for normalization (Saint-Marcoux et al., 2015), and expression values were calculated using the 2(^−ΔΔCt^) method (Livak and Schmittgen, 2001). Gene-specific primers are listed in table S10. Every experiment was performed using three biological replicates, and all expression values were normalized against the average of untreated wild type, which was set to 1.

### Yeast two-hybrid

Yeast two-hybrid assays were performed as described previously (Rizzini et al., 2011). Gateway-based cloning was used to insert the Mp*UVR8.1*, Mp*UVR8.2*, Mp*RUP*, Mp*COP1*, and Mp*SPA* coding sequences downstream of the LexA DNA binding domain (BD) in pBTM116-D9 (Stelzl et al., 2005) or GAL4 activation domain (AD) in pGADT7-GW (Marrocco et al., 2006). The empty vectors were used as negative controls. All the constructs were transformed into the yeast strain L40 using the lithium-acetate protocol (Gietz et al., 2014). For the interaction assays, serial dilutions of transformed yeast colonies were grown under narrowband UV-B (1.5 µmol m^-2^ s^-1^) at 30°C for three days either under a WG305 (+UV-B) or a WG368 (-UV-B control) cut-off filter (Schott Glaswerke). For quantification, transformed yeast colonies were grown under +UV-B or - UV-B control conditions for 16 h. Ten colonies were then combined and resuspended in YPD media, OD_600_ was measured, and cells were spun down and washed. The quantitative interaction assay was carried out with chlorophenol red-β-D-galactopyranoside (CPRG; Roche Applied Science) as substrate, according to the Yeast Protocols Handbook (Clontech, Version PR973283).

### Affinity purification – mass spectrometry (AP–MS)

AP–MS was performed using a Marchantia transgenic line expressing *Pro*_Mp*UVR8*_*:GFP-*Mp*UVR8*, with wild-type Tak-1 and a *Pro*_Mp*UBQ2*_*:mVenus* line as negative controls. Plants were grown for 7 days under continuous white light (50 μmol m⁻² s⁻¹) without UV-B, or under continuous white light (50 μmol m⁻² s⁻¹) supplemented with narrowband UV-B (1.5 μmol m⁻² s⁻¹) followed by a 20 min broadband UV-B exposure (25 μmol m⁻² s⁻¹) immediately prior to harvest. Total protein was extracted and GFP-Trap agarose beads (Chromotek) were used for affinity purifications. Eluted proteins were subjected to LC-MS/MS analysis.

Sample preparation and LC-MS/MS data acquisition: enriched proteins were submitted to an on-bead digestion using trypsin. In brief, dry beads were re-dissolved in 25 µL digestion buffer 1 (50 mM Tris, pH 7.5, 2M urea, 1mM DTT, 5 ng/µL trypsin) and incubated for 30 min at 30 °C in a Thermomixer with 400 rpm. Next, beads were pelleted and the supernatant was transferred to a fresh tube. Digestion buffer 2 (50 mM Tris, pH 7.5, 2M urea, 5 mM CAA) was added to the beads, after mixing the beads were pelleted, the supernatant was collected and combined with the previous one. The combined supernatants were then incubated o/n at 32°C in a Thermomixer with 400 rpm; samples were protected from light during incubation. The digestion was stopped by adding 1 µL TFA and desalted with C18 Empore disk membranes according to the StageTip protocol (Rappsilber et al., 2003).

Dried peptides were re-dissolved in 2% ACN, 0.1% TFA (10 µL) for analysis. Samples were analyzed using an EASY-nLC 1200 (Thermo Fisher) coupled to a Q Exactive Plus mass spectrometer (Thermo Fisher). Peptides were separated on 16 cm frit-less silica emitters (New Objective, 75 µm inner diameter), packed in-house with reversed-phase ReproSil-Pur C18 AQ 1.9 µm resin (Dr. Maisch). Peptides were loaded on the column and eluted for 115 min using a segmented linear gradient of 5% to 95% solvent B (0 min : 5%B; 0-5 min -> 5%B; 5-65 min -> 20%B; 65-90 min ->35%B; 90-100 min -> 55%; 100-105 min ->95%, 105-115 min ->95%) (solvent A 0% ACN, 0.1% FA; solvent B 80% ACN, 0.1%FA) at a flow rate of 300 nL/min. Mass spectra were acquired in data-dependent acquisition mode with a TOP15 method. MS spectra were acquired in the Orbitrap analyzer with a mass range of 300–1750 m/z at a resolution of 70,000 FWHM and a target value of 3×10^6^ ions. Precursors were selected with an isolation window of 1.3 m/z (Q Exactive Plus). HCD fragmentation was performed at a normalized collision energy of 25. MS/MS spectra were acquired with a target value of 10^5^ ions at a resolution of 17,500 FWHM, a maximum injection time (max.) of 55 ms and a fixed first mass of m/z 100. Peptides with a charge of +1, greater than 6, or with unassigned charge state were excluded from fragmentation for MS^2^, dynamic exclusion for 30s prevented repeated selection of precursors.

Data analysis: raw data were processed using MaxQuant software (version 1.6.3.4, http://www.maxquant.org/) (Cox and Mann, 2008) with label-free quantification (LFQ) and iBAQ enabled (Tyanova et al., 2016).

MS/MS spectra were searched by the Andromeda search engine against a combined database containing the sequences from *M. polymorpha* (MpTak_v6.1r2.protein.fasta; https://marchantia.info/download/MpTak_v6.1r2/) and sequences of 248 common contaminant proteins and decoy sequences. Trypsin specificity was required and a maximum of two missed cleavages allowed. Minimal peptide length was set to seven amino acids. Carbamidomethylation of cysteine residues was set as fixed, oxidation of methionine and protein N-terminal acetylation as variable modifications. The match between runs option was enabled. Peptide-spectrum-matches and proteins were retained if they were below a false discovery rate of 1% in both cases. The MS data have been deposited in the ProteomeXchange Consortium via the PRIDE partner repository with the dataset identifier PXD065691.

Statistical analysis of the MaxLFQ values was carried out using Perseus (version 1.5.8.5, http://www.maxquant.org/). Quantified proteins were filtered for reverse hits and hits “identified by site” and MaxLFQ values were log2 transformed. After grouping samples by condition only those proteins were retained for the subsequent analysis that had three valid values in one of the conditions. Missing values were imputed from a normal distribution (1.8 downshift, separately for each column). Significantly enriched proteins were identified using a p-value cutoff of < 0.01 and a log10 fold change threshold of ±1.0 (table S11).

### F_v_/F_m_ measurements

Plates containing 15 gemmalings per genotype were incubated in darkness for at least 5 min, which was sufficient to ensure a steady state under the plant growth conditions. A PSI (Photon Systems Instruments) FluorCam 800 MF was used to measure maximum fluorescence in the dark (F_m_) using saturating pulses of 2000 µmol m⁻² s⁻¹ white light (400–720 nm) for 960 ms, followed by detection pulses of orange-red light (620 nm) lasting 10 µs. The F_v_/F_m_ ratio was calculated using the formula F_v_/F_m_ = (F_m_ – F_o_) / F_m_, where Fm represents the maximal fluorescence, and F_o_ represents the minimal fluorescence in the dark-adapted state.

### Luciferase complementation imaging (LCI) and reporter assays

LCI assays were performed as previously described (Zhou et al., 2018). Site-directed mutagenesis was performed as described previously (Sawano and Miyawaki, 2000) to generate MpUVR8.1^D95N,D106N^ and MpUVR8.2^D95N,D106N^. The pCambia1300–nLUC vector was first digested with BamHI and SalI, while the pCambia1300–cLUC vector was digested with KpnI and SalI. Subsequently, the cDNA fragments of Mp*UVR8.1*, Mp*UVR8.2*, Mp*UVR8.1^D95N,D106N^*, Mp*UVR8.2^D95N,D106N^*, Mp*RUP*, and Mp*COP1* were cloned into the two digested vectors using Gibson assembly. *Agrobacterium tumefaciens* strain GV3101 harboring the indicated combinations of constructs expressing nLUC and cLUC fusion proteins were mixed in a 1:1 ratio and infiltrated into *N. benthamiana* leaves. The plants were kept in the growth chamber for 2 days before being incubated with 1 mM luciferin (D-luciferin sodium salt, Yeasen Biotechnology) for 10 min (in darkness). Luciferase activity was detected using a NightSHADE LB985 In Vivo Plant Imaging System (Berthold).

For *Pro_MpCHS2_:LUC* luciferase reporter assays, gemmae were transferred into 96-well plates containing ½-strength Gamborg’s B5 medium with 1.2% (w/v) agar and grown under at 22°C continuous white light (50 μmol m^-2^ s^-1^) in a CLF GroBank growth chamber for two days. Four-µL of 1 mM luciferin were added to each well and the following day plants were exposed to UV-B for 20 min and luminescence was recorded using a Centro XS LB 960 luminometer.

### Extraction of phenolic compounds, measurements of absorption spectra, and high-performance thin-layer chromatography (HPTLC)

Phenolic compounds were extracted in 100 μL of 80% (v/v) methanol from 50 mg of fresh plant material homogenized using a Silamat S6 (Ivodar Vivadent). Extracts were incubated for 10 min at 70°C on a shaker and centrifuged for 10 min at 12,000 *g* at room temperature (RT). A 40-µL aliquot of clear supernatant from each extract was transferred to a transparent 96-well plate and absorption spectra (200–1000 nm wavelengths) were measured using a TECAN Spark 10M Multimode Microplate Reader. In case absorbance values exceeded the linear range, aliquots were uniformly diluted with extraction buffer.

HPTLC was used to analyze the phenolic compound profile using silica-60 HPTLC glass plates (Sigma-Aldrich), as described previously (Stracke et al., 2010; Leonardelli et al., 2024).

### Microscopy

*Pro*_Mp*UVR8*_*:GFP-*Mp*UVR8* transgenic gemmalings were grown for 7 days under 50 µmol m^−2^s^−1^ white light and irradiated for 20 min with 25 μmol m^-2^ s^-1^ broadband UV-B. Imaging was performed using a Leica Thunder Widefield microscope in both GFP and bright-field modes. Z-stack images were acquired at 2-µm intervals. Image processing and Z-projection were conducted using ImageJ software.

### Transcriptome analysis

Gemmalings were grown for 7 days in 50 μmol m^-2^ s^-1^ white light and exposed or not to supplemental UV-B (1.5 μmol m^-2^ s^-1^) for 3 h. Total RNA was extracted using the RNeasy Plant Mini kit (Qiagen) and Illumina Tru-Seq libraries were generated from cDNA and sequenced using Illumina NovaSeq 100-bp single-end (table S3). Raw reads were trimmed for quality using Trim Galore (v0.6.7; github.com/FelixKrueger/TrimGalore) with “-q 25 --stringency 5” as parameters. Trimmed reads were mapped to the MpTak.v7.1 genome reference using STAR (v2.7.4; (Dobin et al., 2013), followed by read quantification using featureCounts (Subread v2.0.1; (Liao et al., 2019). The raw counts were normalized and differential expression analysis was performed using DESeq2 (Love et al., 2014). Genes with a log_2_ fold change greater than 1 and a p-adj value of less than 0.05 were considered as significantly differentially expressed. Genes with lower log2 fold change values and p-adj < 0.05 were considered when comparing different genotypes in which gene expression changes may occur at varying magnitude (table S4). Euler plots were generated using the eulerr package in R. Heatmaps were generated using the heatmap.2 function in the gplots package in R. The raw RNA-sequencing data has been deposited in the NCBI Gene Expression Omnibus (www.ncbi.nlm.nih.gov/geo) and is accessible through GEO Series accession number GSE296364.

### Untargeted metabolomics

7-day-old Marchantia gemmalings were ground in liquid nitrogen using a mortar and pestle. 50 mg of tissue powder were transferred into a 1.5-mL tube and compounds were extracted with 5 volumes of 80% (v/v) methanol containing 0.1% (v/v) formic acid. After centrifugation, the supernatant was transferred to an HPLC glass vial fitted with a conical insert. A 0.75-µL injection of the extract was performed into ultrahigh performance liquid chromatography coupled to high-resolution mass spectrometry (UHPLC-HRMS) equipped with an electrospray ionization (ESI) source operating in negative (ESI-) ionization mode. Specifically, the system used was an Acquity UPLC I-Class (Waters) interfaced with a Synapt XS QTOF (Waters). The parameters used for untargeted metabolomics were based on (Defossez et al., 2023) with minor adjustments. Briefly, chromatography was done on a Waters Acquity UPLC™ HSS T3 column (100 × 2.1 mm, 1.8 μm) with mobile phases consisting of H2O + 0.05% (v/v) formic acid (solvent A) and acetonitrile + 0.05% (v/v) formic acid (solvent B). A linear gradient from 0 to 100% B in 10.0 min at a flow rate of 0.5 mL/min was applied. MS detection was performed in negative electrospray ionisation using data-dependent acquisition (DDA). The mass range for both MS and MS/MS was 50–1200 Da. Scan times for MS and MS/MS were 0.1 and 0.05 s, respectively. The top 7 precursor ions were selected for MS/MS with an intensity threshold of 15,000 counts per second. An exclusion list of the 150 most intense background ions was generated from a blank sample run just before the plant samples, and a dynamic exclusion of 1.5 s was applied after each MS/MS acquisition. For MS/MS acquisition, a ramped collision energy of 5–40 V (at *m/z* 50) and 20–70 V (at *m/z* 1200) was set. The entire system and method were controlled by MassLynx 4.2 (Waters). Raw profile data were first centroided in MassLynx and further converted to MzML using MSConvert (ProteoWizard). We used MZmine 4.5 for feature detection, deconvolution, and alignment of DDA (data-dependent acquisition) data, with noise thresholds set to 2000 for MS1 and 50 for MS2 (Schmid et al., 2023). The resulting peak table and MS/MS spectral data were exported and annotated using SIRIUS 6.2 (Duhrkop et al., 2019), including CSI:FingerID for structural identification and CANOPUS for the prediction of compound classes, superclasses, and biosynthetic pathways. Taxonomically informed reannotation of molecular features was performed using the Tima R package (Rutz and Allard, 2025).

### Statistical analyses

Statistical analyses were performed using GraphPad Software Prism 10 software (San Diego, California). Statistical significance of the differences between means was determined using one- or two-way analysis of variance (ANOVAs) followed by Tukey’s post hoc honestly significant difference (HSD) test for multiple comparisons. For metabolomics, adjusted p-values were calculated and corrected using the Benjamini-Hochberg in R.

### Accession numbers

Sequence data from this article can be found in the GenBank/EMBL data libraries under accession numbers: Mp2g11590 (Mp*UVR8*), Mp2g18040 (Mp*RUP*), Mp5g12010 (Mp*COP1*), Mp3g25460 (Mp*SPA*), Mp1g16800 (Mp*HY5*), Mp2g07060 (Mp*CHS2*), Mp1g10150 (Mp*PAL1*), Mp6g00020 (Mp*C4H1*), Mp8g05070 (Mp*C4H2*), Mp8g05080 (Mp*C4H3*), Mp1g27780 (Mp*4CL1*), Mp1g05060 (Mp*4CL2*), Mp2g22700 (Mp*CHS6*), Mp5g01410 (Mp*CHIL2*), Mp6g05290 (Mp*CHI*), Mp1g26540 (Mp*F3H*), Mp3g23400 (Mp*EF1α*).

## Supplementary materials

## ACKNOWLEDGEMENTS

We would like to thank Youichi Kondou (Kanto Gakuin University College of Science and Engineering, Yokohama, Japan) for kindly providing the Mp*uvr8-1* mutant and the pMpGWB301-MpUVR8 construct, Félix Rico-Reséndiz (University of Geneva) for sharing protocols and tips regarding Marchantia cultivation and transformation, Yanfei Zhou (The New Zealand Institute for Plant and Food Research Limited, Palmerston North, New Zealand) for precious suggestions, and Anne Harzen (Max-Planck Institute for Plant Breeding Research) for MS sample preparation. The RNA-seq experiments were performed at the iGE3 Genomics Platform of the University of Geneva.

## AUTHOR CONTRIBUTIONS

Y.L., R.P., and R.U. conceived and designed the research; Y.L., R.P., and R.C. performed the experiments; G.G. and E.Def. performed and contributed metabolomic data; S.C.S and H.N. performed and analyzed LC-MS/MS data; J.R. and L.D. contributed Mp*hy5* mutants; Y.L., R.P., E.Dem., and R.U. analyzed the data and wrote the manuscript. All authors reviewed and approved the submitted manuscript.

---

**Figure S1.**
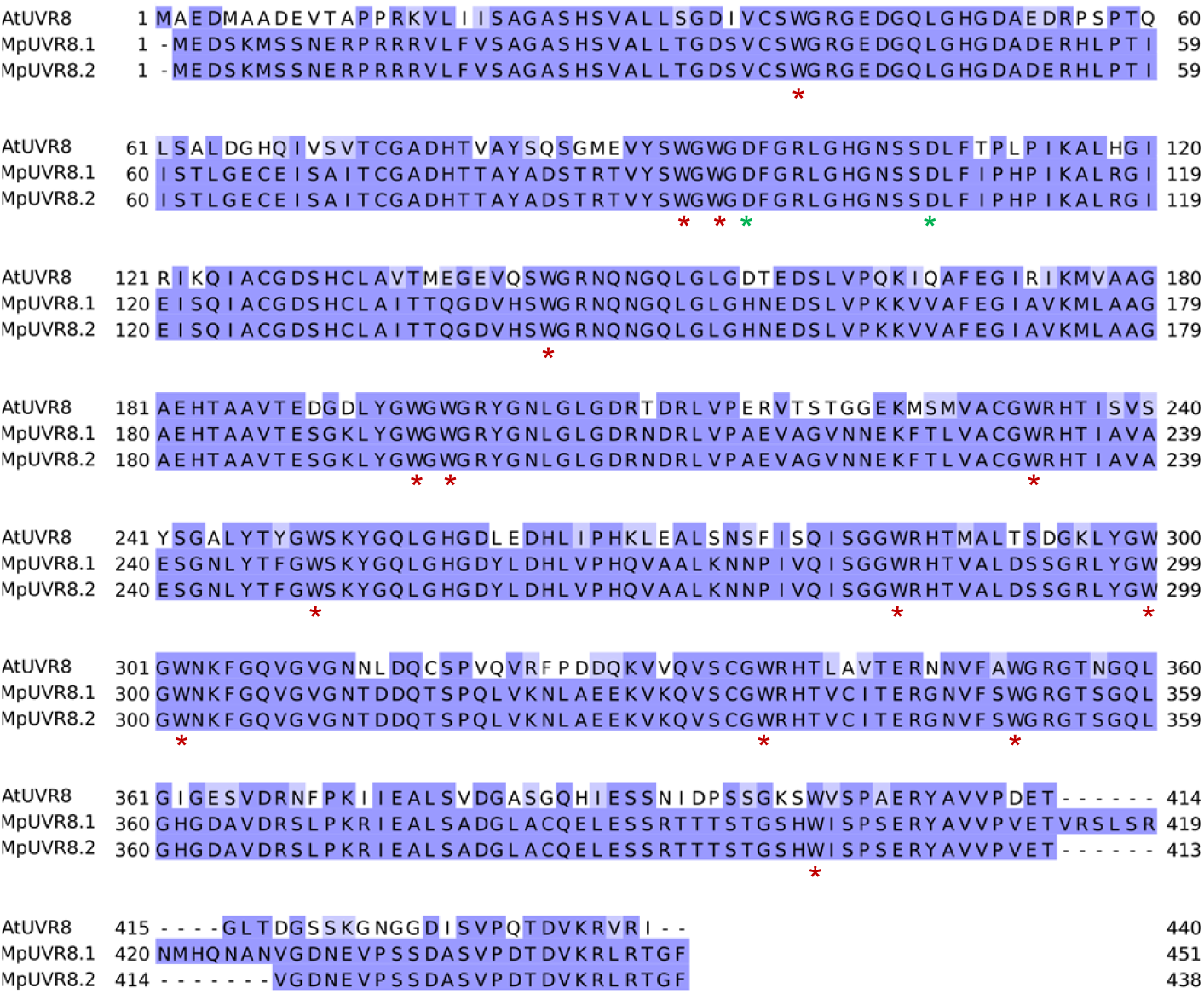
Alignment of Marchantia MpUVR8.1 and MpUVR8.2 with Arabidopsis AtUVR8. Red asterisks (*) highlight conserved tryptophan (W) amino acid residues, including the AtUVR8 key triad tryptophans W233 (W232 in MpUVR8), W285 (W284 in MpUVR8), and W337 (W336 in MpUVR8), within their conserved GWRHT sequence motif. Green asterisks (*) highlight AtUVR8 aspartates D96 (D95 in MpUVR8) and D107 (D106 in MpUVR8).

**Figure S2.**
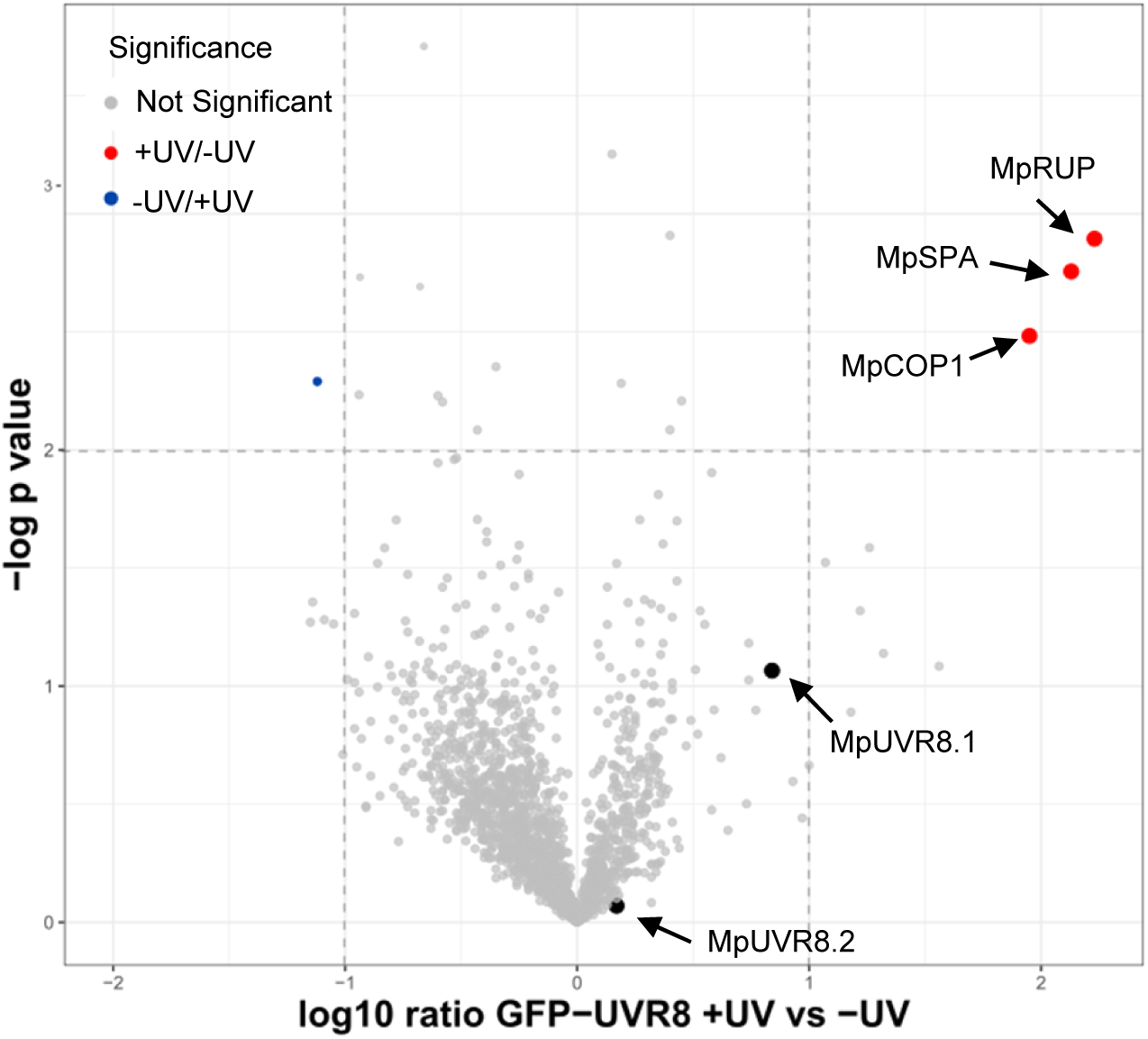
Volcano plot of GFP-MpUVR8 AP-MS under +UV versus -UV conditions. AP–MS was performed using a Marchantia *Pro*_Mp*UVR8*_*:GFP-* Mp*UVR8* (genomic) transgenic line grown for 7 days under continuous white light (50 μmol m⁻² s⁻¹) without UV-B, or under continuous white light supplemented with narrowband UV-B (1.5 μmol m⁻² s⁻¹) followed by a 20 min broadband UV-B exposure (25 μmol m^-2^ s^-2^). Significantly enriched proteins were identified using a p-value cutoff of < 0.01 and a log10 fold change threshold of ±1.0. Grey dots represent non-significant proteins; red dots indicate proteins significantly enriched in +UV compared to –UV; blue dots indicate proteins significantly enriched in –UV compared to +UV. Arrows highlight proteins of interest.

**Figure S3.**
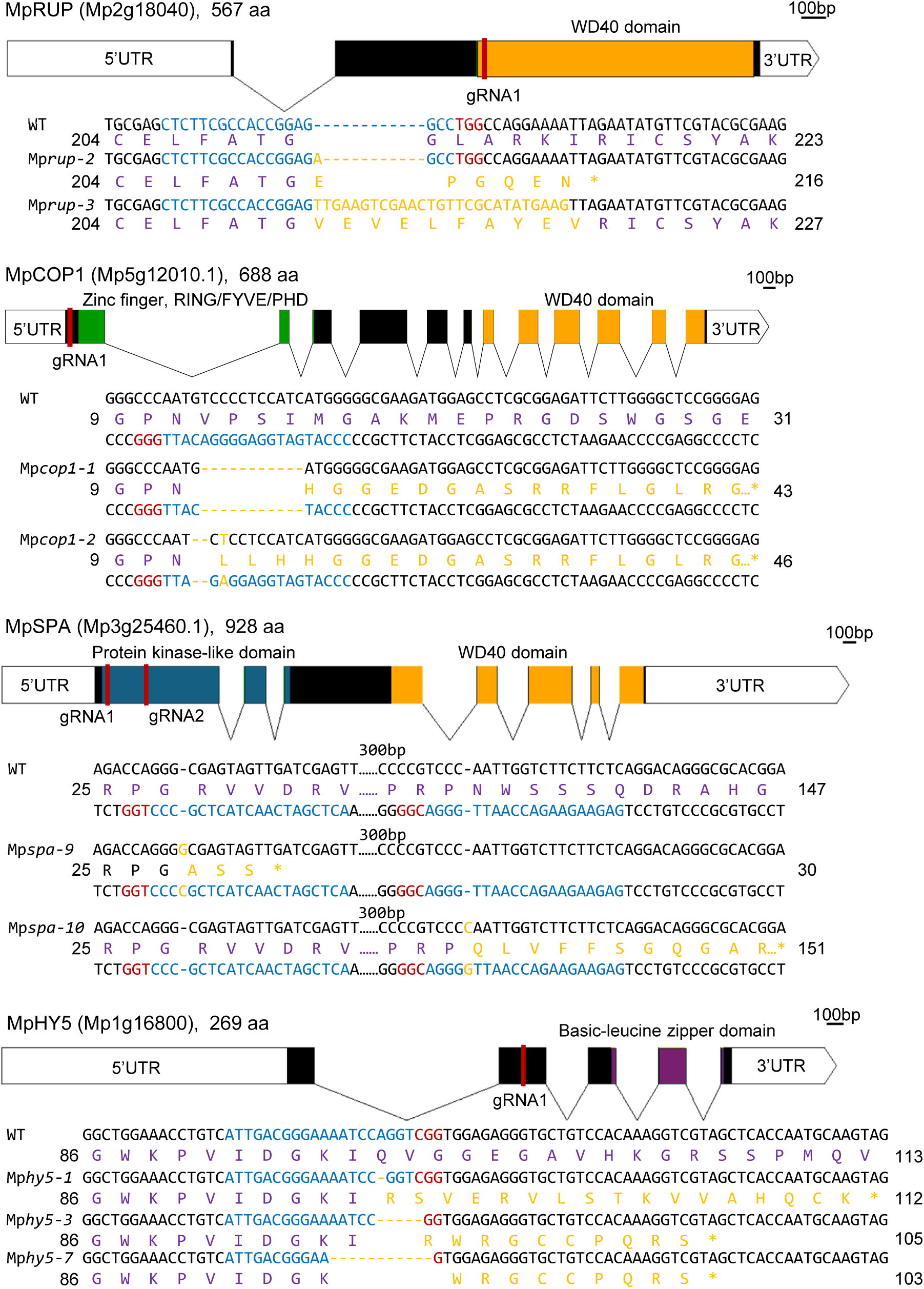
Gene structure and indication of the mutated sites in the corresponding mutants generated by CRISPR/Cas9. Untranslated (5ʹ and 3ʹ UTRs) and coding sequences are highlighted as white and colored boxes, respectively, with specific domains indicated. Introns are represented by lines connecting exons. The positions of the guide RNA (gRNA) (red bar, sequence in blue), protospacer adjacent motif (PAM) sequences (sequence in red), different mutations including resulting amino acid changes (in yellow), translated amino acids of wild-type (in purple) and the number of translated amino acids are also highlighted. The Mp*rup-2* mutant exhibits an insertion of one adenine (A) nucleotide, while Mp*rup-3* displays 27-bp insertion with a 15-bp deletion of endogenous sequence. Mp*cop1-1* and Mp*cop1-2* mutants show an 11-bp deletion and a 2-bp deletion with one cytosine (C) nucleotide substituted by thymine (T), respectively. Mp*spa-9* has an insertion of one guanine (G) nucleotide for gRNA1, whereas Mp*spa-10* exhibits an insertion of one cytosine (C) nucleotide for gRNA2. The Mp*hy5-1* mutant displays a deletion of one adenine (A) nucleotide, whereas Mp*hy5-3* and Mp*hy5-7* mutants show a 5-bp deletion and 11-bp deletion, respectively. All these mutations, except for Mp*rup-3*, result in a frameshift and early STOP codon in the coding sequence.

**Figure S4.**
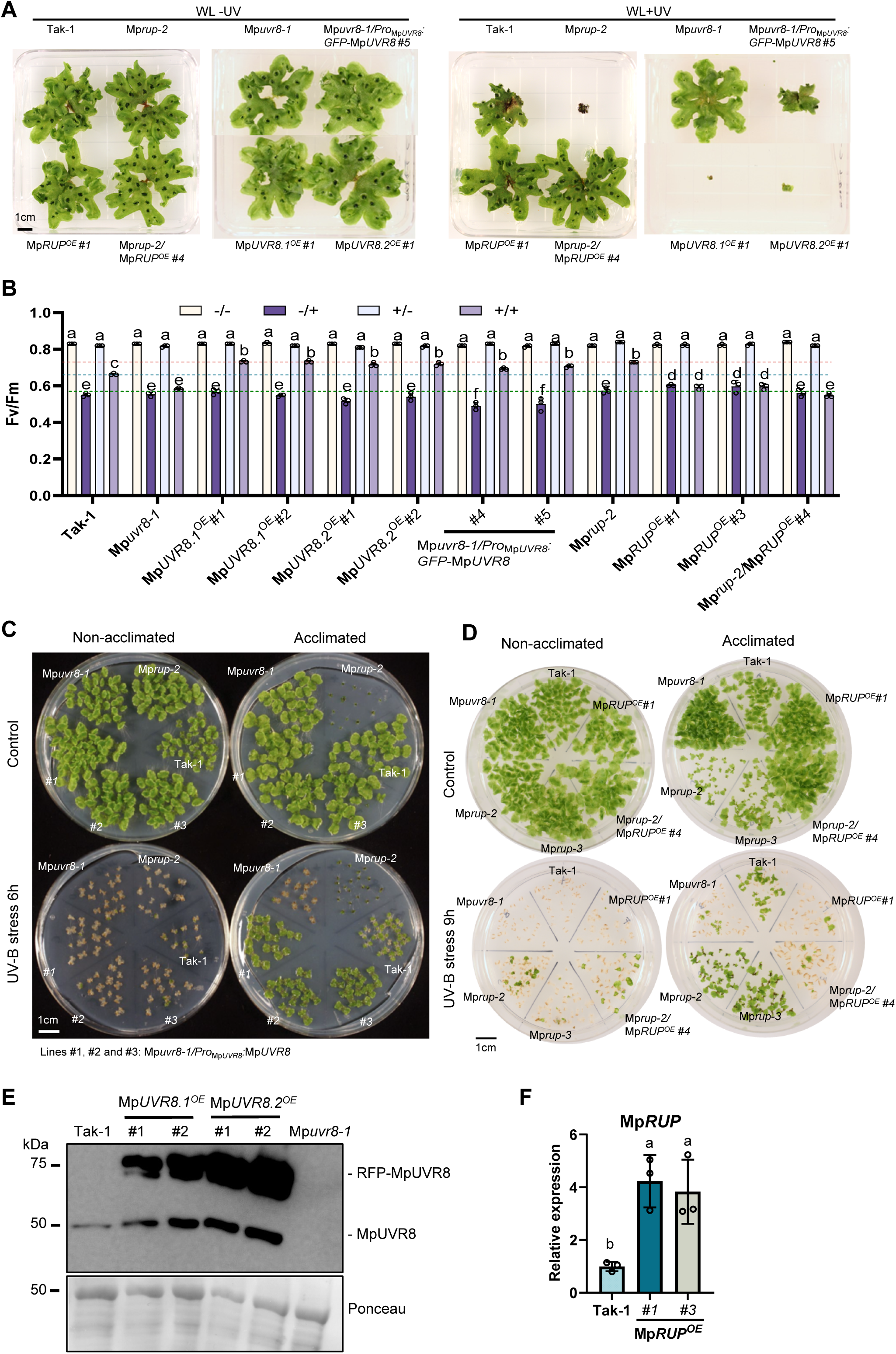
UV-B photomorphogenesis and photoprotection in MpUVR8 and MpRUP transgenic lines. **A)** Growth phenotype of 4-week-old wild-type (Tak-1), Mp*rup-2*, Mp*uvr8-1*, Mp*uvr8-1/Pro*_Mp*UVR8*_*:GFP-*Mp*UVR8* line *#5,* Mp*RUP^OE^ #1*, Mp*rup-2/*Mp*RUP^OE^ #4*, Mp*UVR8.1^OE^ #1*, and Mp*UVR8.2^OE^ #1* plants grown in white light supplemented (WL +UV) or not (WL -UV) with UV-B. Bar = 1 cm. **B)** F_v_/F_m_ measurements in wild-type (Tak-1), Mp*uvr8-1*, Mp*UVR8.1^OE^* lines *#1* and *#2*, Mp*UVR8.2^OE^ #1* and *#2*, Mp*uvr8-1/Pro*_Mp*UVR8*_*:GFP-*Mp*UVR8 #4* and *#5,* Mp*rup-2*, Mp*RUP^OE^ #1* and *#3*, and Mp*rup-2/*Mp*RUP^OE^ #4* grown for 7 days in white light (non-acclimated) or white light supplemented with UV-B (acclimated), without (non-stressed) or with (stressed) additional exposure to broadband UV-B stress for 60 min. -/-, non-acclimated, non-stressed; -/+, non-acclimated, stressed; +/-, acclimated, non-stressed; +/+, acclimated, stressed. Individual data points of biological replicates and means ± SD are shown (*n* = 3). Dashed horizontal lines indicate levels corresponding to acclimated and stressed Tak-1 (blue), Mp*uvr8-1* (green), and Mp*rup-2* (orange) plants. Different letters indicate statistically significant differences between the means (P < 0.01) as determined by two-way ANOVA followed by Tukey’s post hoc honestly significant difference (HSD) test. **C** and **D)** UV-B stress tolerance of indicated plant genotypes grown for 7 days in white light supplemented (acclimated) or not (non-acclimated) with UV-B, then exposed or not to 6 hours of broadband UV-B. Pictures were taken after recovery for another 7 days in white light. Bar = 1 cm. (C) #1, #2 and #3: three independent Mp*uvr8-1/Pro*_Mp*UVR8*_:Mp*UVR8* lines. **E)** Immunoblot analysis of MpUVR8 and RFP-MpUVR8 levels in 14-day-old Tak-1, Mp*UVR8.1^OE^#1*, Mp*UVR8.1^OE^#2*, Mp*UVR8.2^OE^#1*, Mp*UVR8.2^OE^#2*, and Mp*uvr8-1* thallus. Ponceau staining is shown as loading control. Mp*UVR8.1^OE^ =* Tak-1/*Pro*_Mp*EF1α*_:RFP-UVR8.1; Mp*UVR8.2^OE^ =* Tak-1/*Pro*_Mp*EF1α*_:RFP-UVR8.2. **F)** RT-qPCR analysis of Mp*RUP* expression level in 14-day-old Tak-1 and two Mp*RUP^OE^* lines. Individual data points of biological replicates and means ± SD are shown (*n* = 3). Different letters indicate statistically significant differences between the means (P < 0.05) as determined by one-way ANOVA followed by Tukey’s post hoc honestly significant difference (HSD) test.

**Figure S5.**
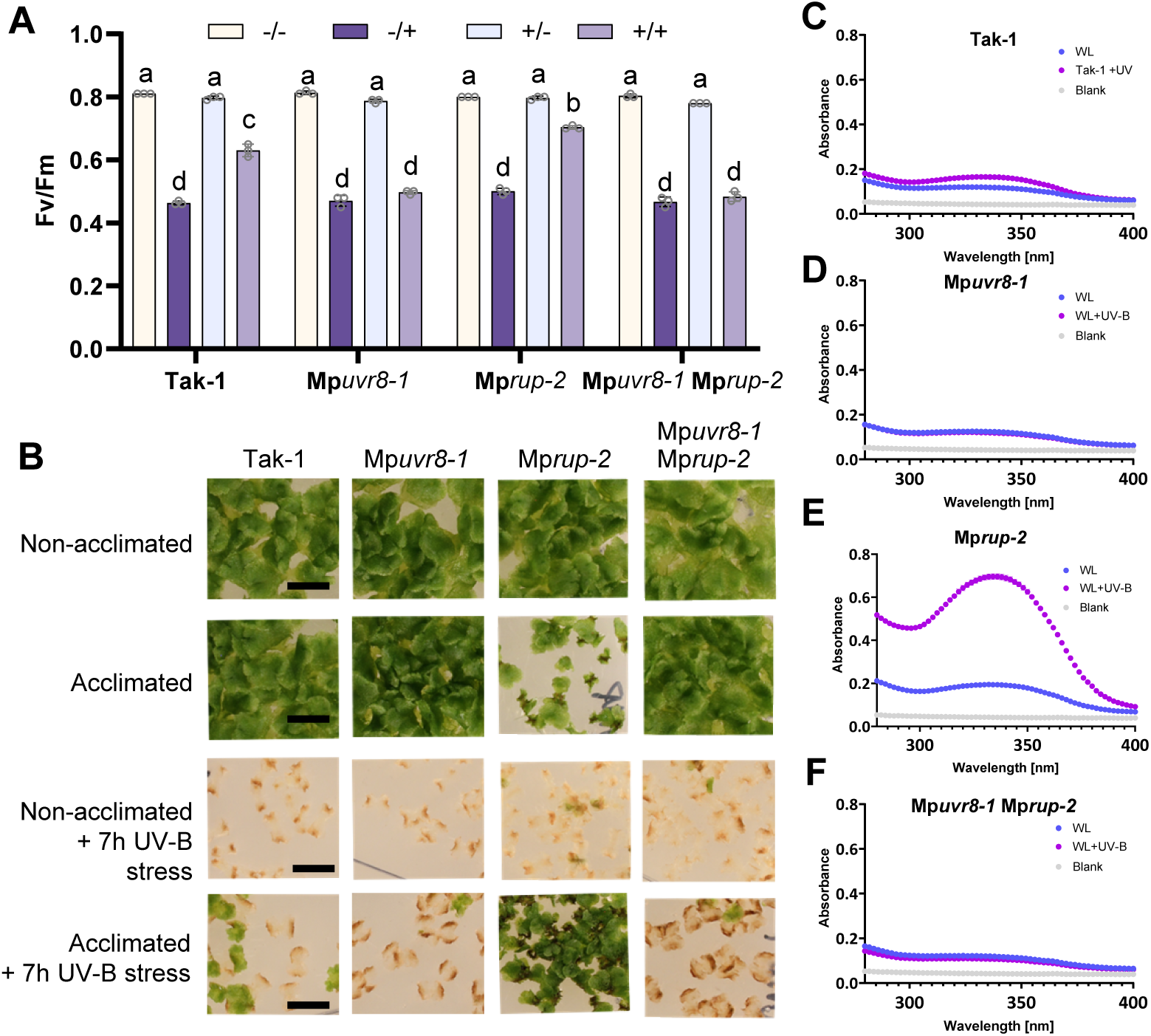
Enhanced photoprotection and UV-B-absorbing capacity of Mp*rup* depends on MpUVR8. **A)** F_v_/F_m_ measurements in wild-type (Tak-1), Mp*uvr8-1*, Mp*rup-2*, and Mp*uvr8-1* Mp*rup-2* grown for 7 days in white light (non-acclimated) or white light supplemented with UV-B (acclimated), without (non-stressed) or with (stressed) additional exposure to broadband UV-B stress for 60 min. -/-, non-acclimated, non-stressed; -/+, non-acclimated, stressed; +/-, acclimated, non-stressed; +/+, acclimated, stressed. Individual data points of biological replicates and means ± SD are shown (*n* = 3). Different letters indicate statistically significant differences between the means (P < 0.01) as determined by two-way ANOVA followed by Tukey’s post hoc honestly significant difference (HSD) test. **B)** UV-B stress resistance in Tak-1, Mp*uvr8-1*, Mp*rup-2*, and Mp*uvr8-1* Mp*rup-2* plants grown for 7 days in white light supplemented (acclimated) or not (non-acclimated) with UV-B, then exposed or not to 7 hours of broadband UV-B. Pictures were taken after recovery for another 7 days in white light. Bars = 1 cm. **C-F)** Absorbance spectra (280–400 nm) of methanolic extracts from 9-day-old Tak-1 (C), Mp*uvr8-1* (D), Mp*rup-2* (E), and Mp*uvr8-1* Mp*rup-2* (F) gemmalings grown for 7 days in white light and exposed (WL+UV-B, purple lines) or not (WL, blue lines) to supplemental UV-B (1.5 μmol m^-2^ s^-1^) for 2 days. The gray line shows the absorbance of the buffer (blank). Values of independent measurements and means ± SEM are shown (*n* = 3).

**Figure S6.**
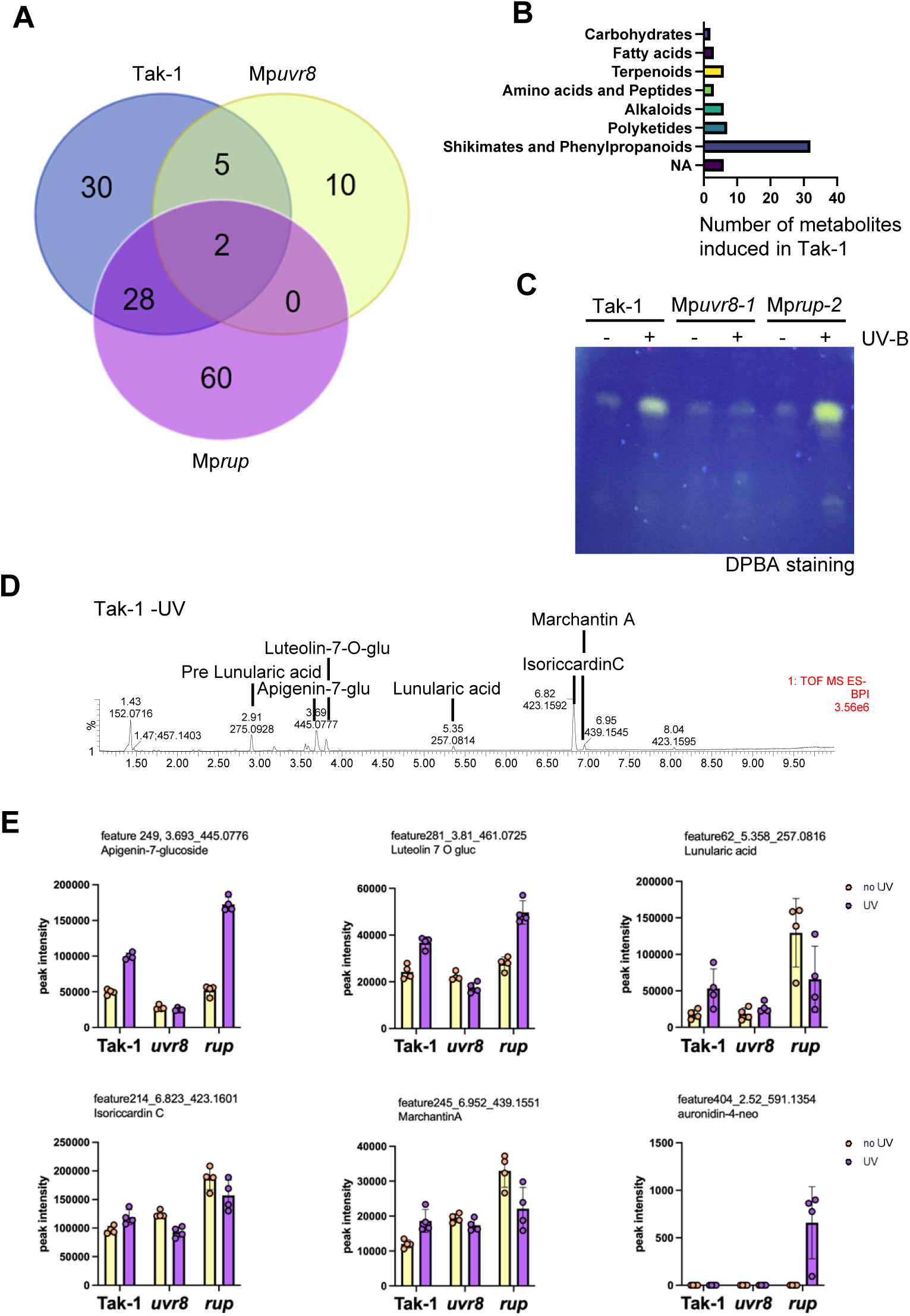
MpUVR8-dependent regulation of metabolite accumulation upon UV-B exposure. **A)** Venn diagram showing overlaps between upregulated metabolites (FC> or =1.5, p-adj < or =0.05) upon exposure to 2 days UV-B in 9-day-old wild-type (Tak-1), Mp*uvr8-1* (Mp*uvr8*), and Mp*rup-2* (Mp*rup*) gemmalings. **B)** Distribution of the number of compounds induced under UV-B in wild type (FC> or =1.5, p-adj < or =0.05), according to their chemical classes. Corresponding metabolite lists: table S7. **C)** High-performance thin-layer chromatography (HPTLC) analysis of flavonol glycoside levels in 7-d-old gemmalings of wild type (Tak-1), Mp*uvr8-1*, and Mp*rup-2* grown for 7 days in white light and exposed, or not, to supplemental UV-B for 2 days. DPBA, diphenylboric acid 2-aminoethylester. **D** and **E)** Metabolomic analysis of 9-day-old plants of Tak-1, Mp*uvr8-1*, and Mp*rup-2* grown for 7 days in white light and exposed (+) or not (-) to supplemental UV-B (1.5 μmol m^-2^ s^-1^) for 2 days (*n* = 5, 12 plants per replicate). (D) UHPLC-MS chromatogram from -UV-B Tak-1 sample. Metabolites corresponding to major peaks are annotated based on their accurate mass and MS2 spectra. (E) Relative quantifications of selected compounds in Tak-1, Mp*uvr8-1* (*uvr8*), and Mp*rup-2* (*rup*) methanolic extracts from plants exposed to UV-B (purple, right bar, UV) or not (yellow, left bar, no UV). Individual data points of biological replicates and means ± SD are shown (*n* = 4).

**Figure S7.**
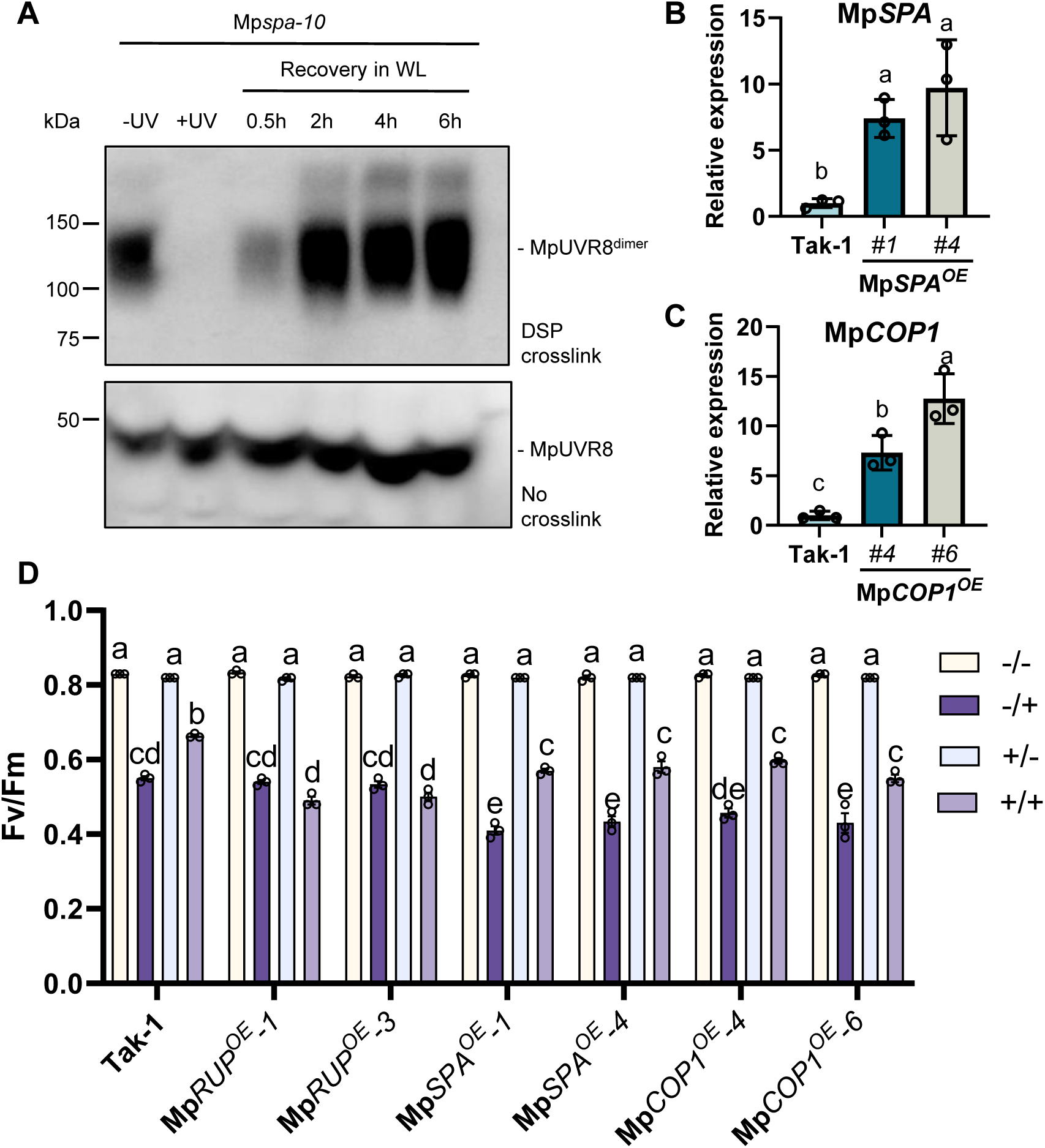
MpSPA does not affect MpUVR8 redimerization, but MpCOP1 and MpSPA overexpression reduces basal UV-B tolerance. **A)** Immunoblot analysis of MpUVR8 levels in 7-day-old Mp*spa-10* gemmalings irradiated for 20 min with broadband UV-B before recovery in white light (WL) for the indicated times. Top: MpUVR8 dimer levels in DSP-crosslinked protein extracts. Bottom: total MpUVR8 levels in non-crosslinked extracts. **B)** RT-qPCR analysis of Mp*SPA* expression level in 14-day-old plants of Tak-1 and Mp*SPA^OE^* lines *#1* and *#4*. Individual data points of biological replicates and means ± SD are shown (*n* = 3). Letters indicate statistically significant differences between the means (P < 0.05) as determined by one-way ANOVA followed by Tukey’s post hoc honestly significant difference (HSD) test. **C)** RT-qPCR analysis of Mp*COP1* expression level in 14-day-old plants of Tak-1, Mp*COP1^OE^* lines *#4* and *#6*. Individual data points of biological replicates and means ± SD are shown (*n* = 3). Letters indicate statistically significant differences between the means (P < 0.05) as determined by one-way ANOVA followed by Tukey’s post hoc honestly significant difference (HSD) test. **D)** F_v_/F_m_ measurements in wild-type (Tak-1), Mp*RUP^OE^* lines *#1* and *#3*, Mp*SPA^OE^ #1* and *#4*, and Mp*COP1^OE^ #4* and *#6* grown for 7 days in white light (non-acclimated) or white light supplemented with UV-B (acclimated), without (not stressed) or with (stressed) additional exposure to broadband UV-B stress for 60 min. -/-, not acclimated, not stressed; -/+, not acclimated, stressed; +/-, acclimated, not stressed; +/+, acclimated, stressed. Individual data points of biological replicates and means ± SD are shown (*n* = 3). Letters indicate statistically significant differences between the means (P < 0.01) as determined by two-way ANOVA followed by Tukey’s post hoc honestly significant difference (HSD) test.

**Figure S8.**
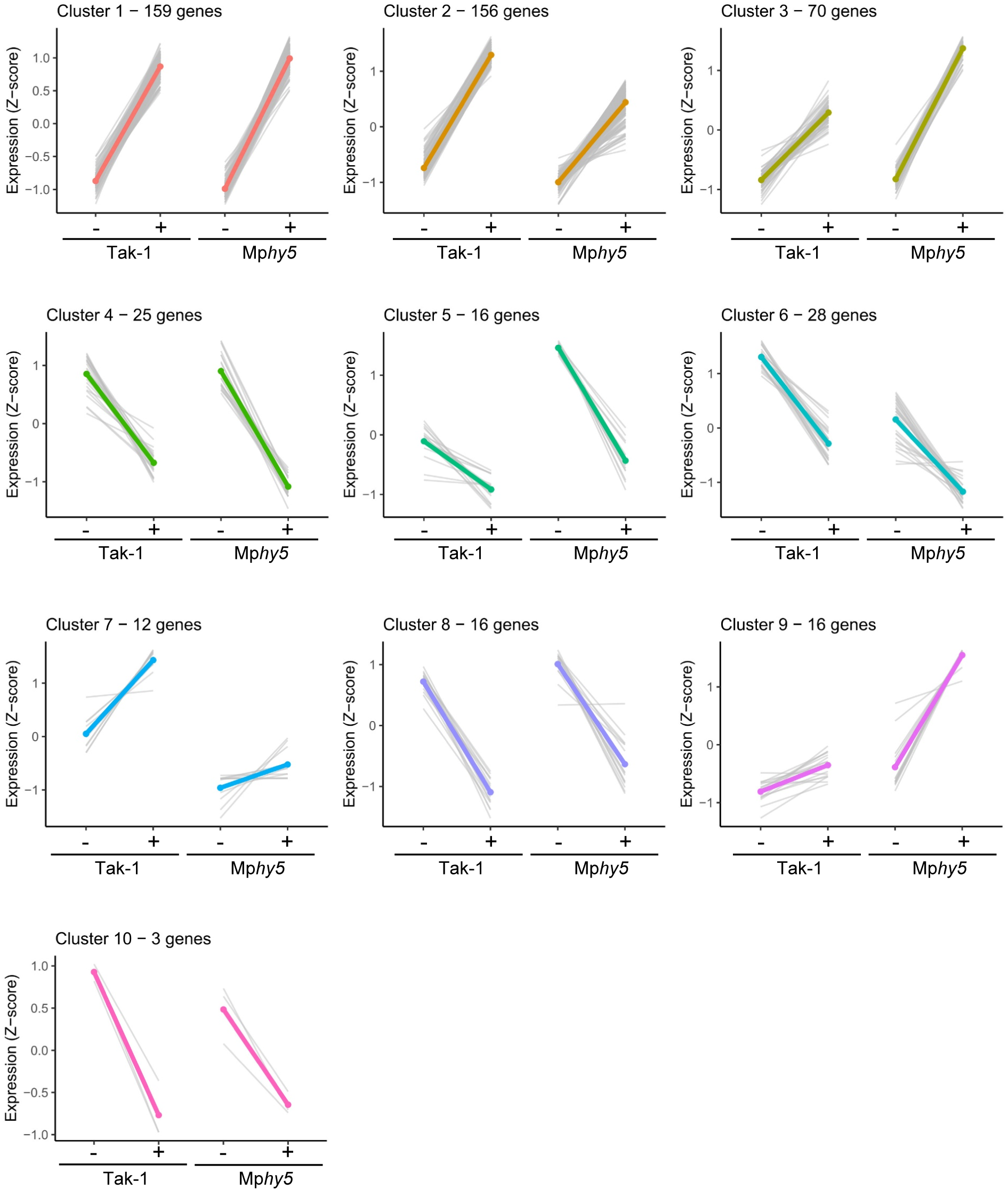
MpHY5 plays a minor role in the regulation of UV-B-induced gene expression changes. Gene expression profiles in clusters identified by hierarchical clustering. Z-scores for normalized read count values for each gene (gray lines) and for the cluster average (colored line) are shown.

**Figure S9.**
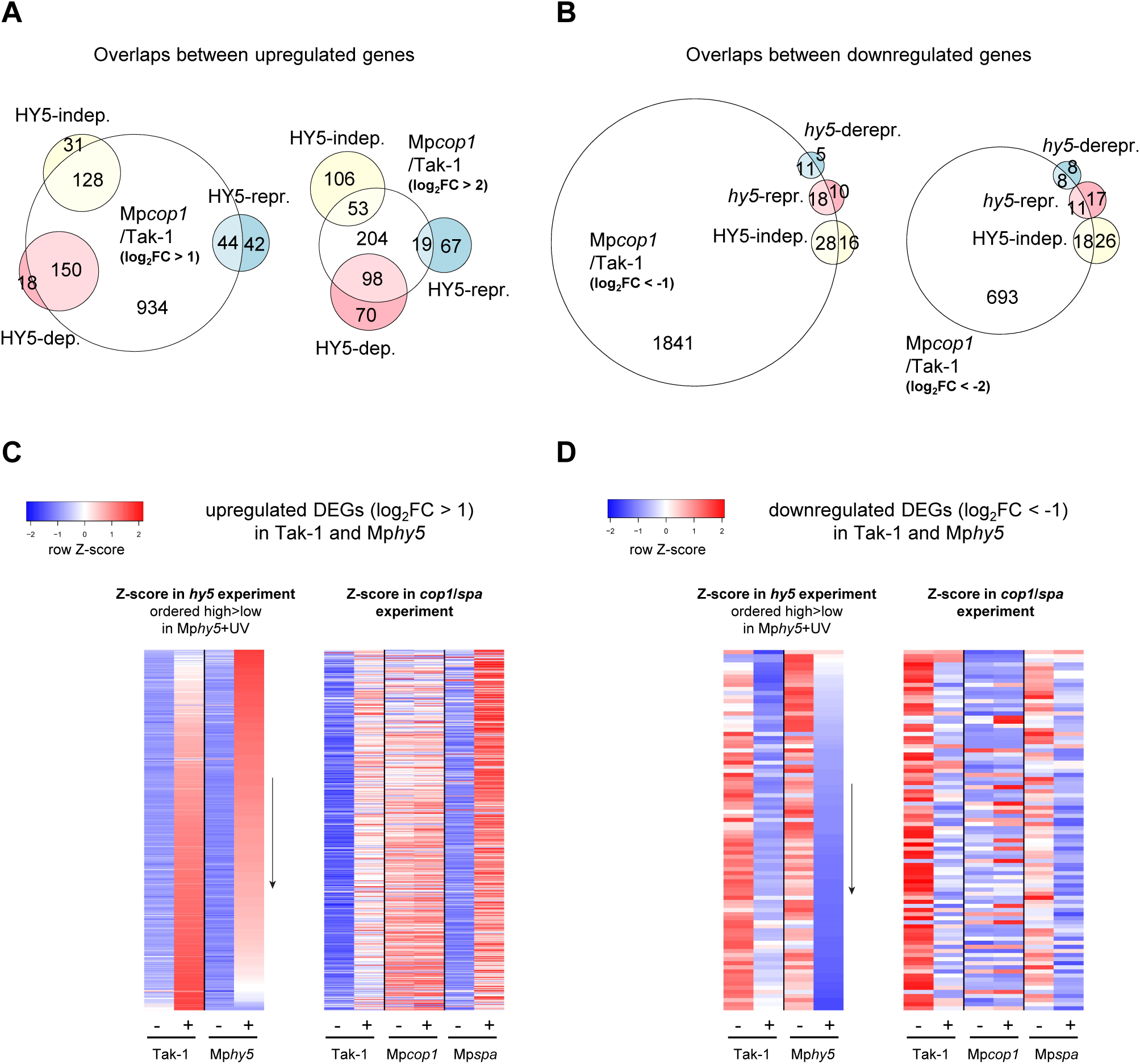
MpHY5 and MpCOP1 regulate UV-B-induced gene expression in an antagonist manner. **A)** Euler plots showing overlaps between upregulated DEGs (p-adj < 0.05) in Mp*cop1*/Tak-1 in white light (left: log_2_FC > 1; right: log_2_FC > 2) and genes that are induced after 3h of UV-B. UV-B-upregulated genes are separated into 3 categories: genes that are induced in a MpHY5-independent manner (“HY5-indep.”, fig. S7 cluster 1), genes that are dependent on MpHY5 for their full induction or hypoinduced in Mp*hy5* (“HY5-dep.”, fig. S7 clusters 2 and 7) and genes that are hyperinduced in Mp*hy5* or whose induction is repressed by HY5 (“HY5-repr.”, fig. S7 clusters 3 and 9). Corresponding gene lists: table S8a. **B)** Euler plots showing overlaps between downregulated DEGs (p-adj < 0.05) in Mp*cop1*/Tak-1 in white light (left: log_2_FC < -1; right: log_2_FC < -2) and genes that are repressed after 3h of UV-B. UV-B-downregulated genes are separated into 3 categories: genes that are repressed in a MpHY5-independent manner (“HY5-indep.”, fig. S7 clusters 4, 8 and 10), genes that are somewhat constitutively repressed in Mp*hy5* (“*hy5*-repr.”, fig. S7 cluster 6) and genes that are derepressed in white light-grown Mp*hy5* (“*hy5*-derepr.”, fig. S7 cluster 5). Corresponding gene lists: table S8b. **C)** Heatmap of all genes upregulated (log_2_FC > 1 and p-adj < 0.05) by UV-B in Tak-1 and/or Mp*hy5*. Red and blue colors indicate normalized read count Z-scores in RNA-seq experiments including Tak-1 and Mp*hy5* (left) and Tak-1, Mp*cop1* and Mp*spa* (right). Z-scores were calculated separately for the two experiments and genes were ordered in a decreasing order according to the Z-score in Mp*hy5*+UV (arrow). **D)** Heatmap of all genes downregulated (log_2_FC < -1 and p-adj < 0.05) by UV-B in Tak-1 and/or Mp*hy5*. Red and blue colors indicate normalized read count Z-scores in RNA-seq experiments including Tak-1 and Mp*hy5* (left) and Tak-1, Mp*cop1* and Mp*spa* (right). Z-scores were calculated separately for the two experiments and genes were ordered in a decreasing order according to the Z-score in Mp*hy5*+UV (arrow).

**Figure S10.**
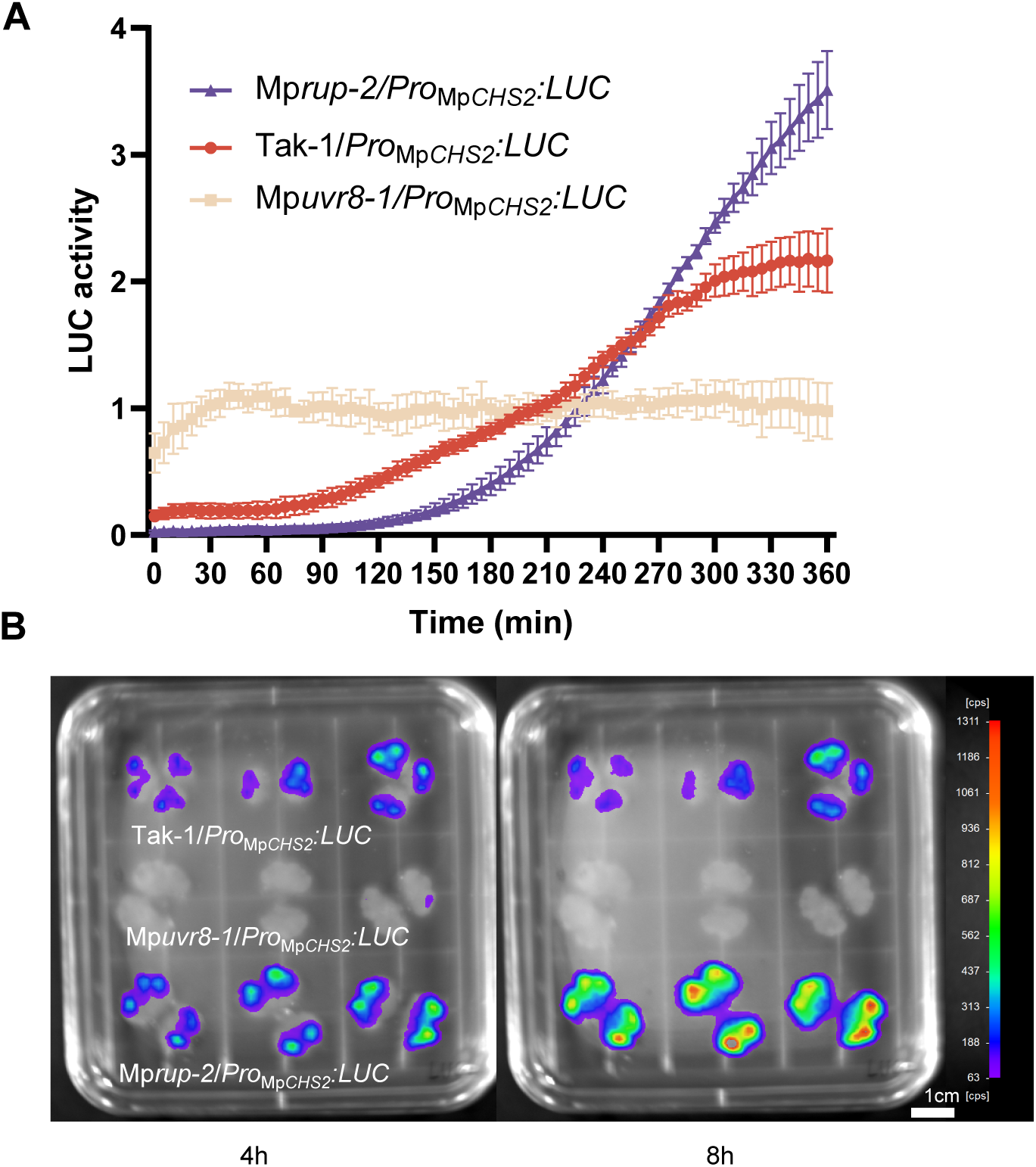
LUC activity of a *Pro*_Mp*CHS2*_*:LUC* reporter in Tak-1, Mp*uvr8-1*, and Mp*rup-2* backgrounds upon UV-B exposure. **A)** The counts per second (cps) of luciferase (LUC) activity were normalized to average expression level measured in 3-day-old gemmalings of the following lines: Tak-1/*Pro*_Mp*CHS2*_*:LUC*, Mp*uvr8-1*/*Pro*_Mp*CHS2*_*:LUC*, and Mp*rup-2*/*Pro*_Mp*CHS2*_*:LUC.* These gemmalings were grown in 96-well plates under white light, and pre-sprayed with luciferin 24 h before a 20-min exposure to broadband UV-B. The LUC activity was monitored for up to 360 min after the UV-B induction. Values of independent measurements and means ± SD are shown (*n* = 8). **B)** Analysis of LUC activity in 14-day-old thalli of Tak-1/*Pro*_Mp*CHS2*_*:LUC*, Mp*uvr8-1*/*Pro*_Mp*CHS2*_*:LUC*, and Mp*rup-2*/*Pro*_Mp*CHS2*_*:LUC* lines followed by 20 min of broadband UV-B exposure at 4 h (left) and 8 h (right).

