## Supplemental Table S10 for "Conservation and divergence of UVR8-COP1/SPA-HY5 signaling in UV-B responses of *Marchantia polymorpha*"

**Table S10. Primer list**

| Purpose | Name | Sequence (5’-3’) |
| --- | --- | --- |
| Cloning | MpSPA_attB1_Fw | GGGGACAAGTTTGTACAAAAAAGCAGGCTTAATGATTAATGGAGGGTCGGAAG |
|  | MpSPA_attB2_stop_Rv | GGGGACCACTTTGTACAAGAAAGCTGGGTACTACACCATTTCCAAAACTTTGAT |
|  | MpCOP1_attB1_Fw | GGGGACAAGTTTGTACAAAAAAGCAGGCTTAATGGAGGGTACTGAGGGCTC |
|  | MpCOP1_attB2_stop_Rv | GGGGACCACTTTGTACAAGAAAGCTGGGTATCAAGGAGCAAGCACAAGCACT |
|  | MpRUP_attB1_Fw | GGGGACAAGTTTGTACAAAAAAGCAGGCTTAATGAAAATGGGAAGCGGAGATG |
|  | MpRUP_attB2_stop_Rv | GGGGACCACTTTGTACAAGAAAGCTGGGTATCAAATAGAATCTACTTGGTGTGC |
|  | MpUVR8_attB1_Fw | GGGGACAAGTTTGTACAAAAAAGCAGGCTTCATGGAAGACTCGAAGATGTCGAG |
|  | MpUVR8_attB2_stop_Rv | GGGGACCACTTTGTACAAGAAAGCTGGGTCTTAAAAGCCCGTCCGTAGCC |
|  | ProMpCHS2-3k_GW_Fw | GGGGACAAGTTTGTACAAAAAAGCAGGCTTCATTATCTGGATACACACACGTTTCAGG |
|  | ProMpCHS2-3k_GW_Rv | GGGGACCACTTTGTACAAGAAAGCTGGGTCCGTTGCAGCTGAGGAAGAGC |
|  | cLUC-Mprup-Fw | CGTCCCGGGGCGGTACCATGAAAATGGGAAGCGG |
|  | cLUC-Mprup-Rv | ACGAACGAAAGCTCTGCAGGTCAAATAGAATCTACTTG |
|  | Mprup- nLUC_Fw | ACGAGCTCGGTACCCGGATGAAAATGGGAAGCGGA |
|  | Mprup-nLUC_Rv | GGACGCGTACGAGATCTGGCAAATAGAATCTACTTGGT |
|  | cLUC-MpUVR8-Fw | CGTCCCGGGGCGGTACCATGGAAGACTCGAAGAT |
|  | cLUC-MpUVR8-Rv | ACGAACGAAAGCTCTGCAGGTTAAAAGCCCGTCCGTAG |
|  | MpUVR8-nLUC_Fw | ACGAGCTCGGTACCCGGATGGAAGACTCGAAGATG |
|  | MpUVR8-nLUC_Rv | GGACGCGTACGAGATCTGGTAAAAGCCCGTCCGTAGC |
|  | cLUC-MpCOP1-Fw | CGTCCCGGGGCGGTACCATGGAGGGTACTGAGGG |
|  | cLUC-MpCOP1-Rv | ACGAACGAAAGCTCTGCAGGTCAAGGAGCAAGCACAAG |
|  | MpCOP1-nLUC_Fw | ACGAGCTCGGTACCCGGATGGAGGGTACTGAGGGC |
|  | MpCOP1-nLUC_Rv | GGACGCGTACGAGATCTGGCAAGGAGCAAGCACAAGC |
|  | MpUVR8genomic_GFP_1_Fw | TTTAATCATTGGAATGTGTTAACACTTTTGCTTTGGCAAAGTG |
|  | MpUVR8genomic_GFP_1_Rv | AGTTCTTCTCCTTTACTCATTCCGACAGCTGAAATATTGAAG |
|  | MpUVR8genomic_GFP_2_Fw | TCAATATTTCAGCTGTCGGAATGGTGAGCAAGGGCGAGGAG |
|  | MpUVR8genomic_GFP_2_Rv | CCGCCGAAACAAATAGCATCGGCGTGGTCTCTCGTTACTCGACATCTTCGAGTCTTCCTTGTACAGCTCGTCCAT |
|  | MpUVR8_D95ND106N_Fw | AGCTGGGGCTGGGGAAACTTTGGAAGACTTGGGCATGGTAACTCTAGTAATCTTTTCATACCCC |
| Crispr | MpRUP_gR2_Fw | CTCGGCTCTTCGCCACCGGAGGCC |
|  | MpRUP_gR2_Rv | AAACGGCCTCCGGTGGCGAAGAGC |
|  | MpCOP1_gR2_Fw | CTCGCCCATGATGGAGGGGACATT |
|  | MpCOP1_gR2_Rv | AAACAATGTCCCCTCCATCATGGG |
|  | MpSPA_gR1_Fw | CTCGACTCGATCAACTACTCGCCC |
|  | MpSPA_gR1_Rv | AAACGGGCGAGTAGTTGATCGAGT |
|  | MpSPA_gR2_Fw | CTCGGAGAAGAAGACCAATTGGGA |
|  | MpSPA_gR2_Rv | AAACTCCCAATTGGTCTTCTTCTC |
|  | MpHY5_gRNA2_F | TCGATTGACGGGAAAATCCAGGTGT |
|  | MpHY5_gRNA2_R | AAAACACCTGGATTTTCCCGTCAAT |
| Geno-typing | Mprup_1&2_seq_Fw | AGCAAGAGACCTCGAACAGG |
|  | Mprup_1&2_seq_Rv | TCACTGTGTGCTTGTGCTCG |
|  | Mpcop1_2_seq_Fw | AAGGAAGCGAGGACCAGCAG |
|  | Mpcop1_2_seq_Rv | GAGACGGATCAGATTTGTGTGG |
|  | Mpspa_1&2_seq_Fw | ACTCTCGTGCTTCATCTGCG |
|  | Mpspa_1&2_seq_Rv | GGCATCCTCGAAACCTTCAG |
|  | MpHY5_F | TCACAGCAGAGATCGAATCC |
|  | MpHY5_R | TGAAATCGCCAACTTAGAGAAC |
| RT-qPCR | MpEF1α_qPCR_Fw | AAGCCGTCGAAAAGAAGGAG |
|  | MpEF1α_qPCR_Rv | TTCAGGATCGTCCGTTATCC |
|  | MpCOP1_qPCR_Fw | CTCATTTCATCAGTGCGGTATG |
|  | MpCOP1_qPCR_Rv | GCAAGCACAAGCACTTTAATTG |
|  | MpRUP_qPCR_Fw | CAGGCACACCAAGTAGATTCTA |
|  | MpRUP_qPCR_Rv | CGACCTCGAATAAATAGTCCGA |
|  | MpSPA_qPCR_Fw | GATCGTTTCGAGTCAATTAGCC |
|  | MpSPA_qPCR_Rv | CTGTCCTGAGAAGAAGACCAAT |
